# Structural order emerges without mechanical order inside living cells

**DOI:** 10.64898/2026.10.15.759452

**Authors:** Mohammad Amin Eskandari, Susanne Ulmer, Doralycia Carranza Zuñiga, Sascha Lambert, Saheli Dey, Claudia Marcelli, Houra Sarparandeh, Ruben Haag, Dorian Marx, Mattias Luber, Bart Eduard Vos, Bernhard Schmitzer, Sarah Köster, Timo Betz

**Affiliations:** Third Institute of Physics - Biophysics, University of Göttingen, Göttingen, Germany; Institute of Computer Science, University of Göttingen, Göttingen, Germany; National Institute of Astrophysics, Optics and Electronics, San Andrés Cholula, México; Institute for the Dynamics of Complex Systems, University of Göttingen, Göttingen, Germany; Institute for X-ray Physics, University of Göttingen, Göttingen, Germany; Institute for Microbiology and Genetics, University of Göttingen, Göttingen, Germany; Cluster of Excellence ’Multiscale Bioimaging: From Molecular Machines to Networks of Excitable Cells’ (MBExC), University of Göttingen, Göttingen, Germany

## Abstract

Intracellular organization is vital for cell function, and disruption of organelle positioning is connected to severe diseases, such as cancer. Given the small length scale of cells and the importance of Brownian motion, maintaining intracellular organization requires mechanical interactions to remain stable. Here, we test the common view that intracellular organization is reflected in intracellular mechanical properties. Using the recently introduced active mechanical fingerprint, we quantify intracellular mechanical properties and active forces to create mechanical maps of patterned RPE1 cells. The organization of actin, microtubules, the Golgi apparatus, the endoplasmic reticulum, and the nucleus reveals pronounced structural polarization across the cell population. To overcome limitations of simple averages, we use a Wasserstein barycenter to calculate an optimal intracellular organization that explains all measurements with minimal cellular distortion. To our surprise, and in contrast to previous reports, we find neither a statistically significant spatial mechanical pattern nor any relevant correlation between the tested structures and intracellular mechanical properties. While there is clear local mechanical anisotropy, the large heterogeneity of mechanical values outweighs any intracellular mechanical gradient. This is further supported by large cell-to-cell variability. Thus, despite having the same genome, cells function properly while their intracellular mechanical properties vary over two orders of magnitude.

## 1 Introduction

The mechanical properties of living cells are crucial for regulating biological functions, such as differentiation [1], migration [2, 3] and division [4, 5]. Across many of these processes, cells establish distinct structural polarity, characterized by front-rear, apical-basal, or anteriorposterior organization of cell compartments [6]. Consequently, the internal structure of the living cell is fundamentally heterogeneous [7, 8] and anisotropic [9].

Prior studies have established that from a biomechanical perspective, living cells cannot be treated as homogeneous and isotropic viscoelastic materials [10]. Instead, the cytoplasm behaves as a complicated material whose mechanical properties are governed by intracellular constituents and subject to spatiotemporal dynamics [11]. Specifically, distinct cytoplasmic regions display different mechanical properties [12]. For instance, the diffusion coefficient which reflects the mechanical properties via generalized Stokes-Einstein [13, 14], exhibits extreme heterogeneity, varying up to 400-fold across the cytoplasm [15]. Furthermore, when geometric confinement induces morphological anisotropy by making cells longer along one axis than the other, the cytoplasm exhibits higher stiffness along that long axis. [16].

Growing evidence also suggests that the mechanical properties and the cell polarity are related [17]. For example, when cells are confined on two-dimensional adhesive micropatterns, both the complex shear modulus and vimentin filament density exhibit radial and front-rear polarity [18]. Moreover, FRET-based sensors have revealed that intracellular tension, strongly correlates with the actin stress fibers [19].

Despite these insights, the heterogeneity and anisotropy of the mechanical properties and how these properties correlate with cell polarity remain largely unaddressed. To resolve this question, we geometrically constrained retinal pigment epithelial (RPE1) cells using adhesive micropatterns to induce a highly reproducible and well-defined structural polarity. RPE1 cells were chosen as they provide a high contractility that improves the overall reproducibility of intracellular organization on patterns. To construct faithful average cell templates for major cytoplasmic components we developed a novel framework that uses optimal transport to reduce the blurring that is reminiscent of the commonly used averaging. Next, we combined this template with active and passive optical tweezers microrheology to systematically measure the mechanical fingerprint [20] mechanical heterogeneity and anisotropy relative to the average cell template.

While this approach confirms a reproducible structural organization, reflected by distinct spatial localization of organelle and cytoskeletal networks across the micropattern, we demonstrate that the cytoplasm is locally (on the probe level) heterogeneous and anisotropic, leaving micron-sized probes to experience a non-uniform and direction-dependent environment. In contrast to previous reports of global (cell level) mechanical gradients matching organelle distributions [18], mapping our local mechanical properties onto the standardized structural template reveals a flat, uniform global mechanical baseline. Ultimately, while underlying organelles and cytoskeletal networks are highly organized, local microstructural heterogeneities leads to a substantial cell-to-cell variability, which obscures global mechanical gradients.

## 2 Standard Framework For Intracellular Mechanical Mapping

The intracellular environment is characterized by structural heterogeneity and anisotropy. An example of a typical actin staining of an RPE1 cell is shown in Fig. 1a. Focusing on three different regions, we found markedly different actin structures such as a branched network in region 1, and thick stress fibers assemblies in regions 2 and 3. However, whereas the fibers in region 2 are highly aligned, region 3 displays a much less aligned organization. Consequently, one would expect different mechanical properties of the intracellular environment as a function of the actin structures as found in the different regions [21, 22]. Moreover, when probing the cytoplasm in the vicinity of stress fibers, the local mechanical properties are expected to vary depending on whether the measurement is aligned in parallel or perpendicularly to the fiber axis [23, 24]. Because the stress fibers in region 2 are oriented differently than those in region 3, their respective mechanical properties should exhibit a distinct directional dependency that corresponds directly to the local alignment of the cytoskeletal filaments. When taking all other intracellular structures into account, the system becomes extremely complex [25, 26]. This raises the question of whether this structural heterogeneity and anisotropy are reflected in the cell’s mechanical properties and activity.

**Figure 1:**
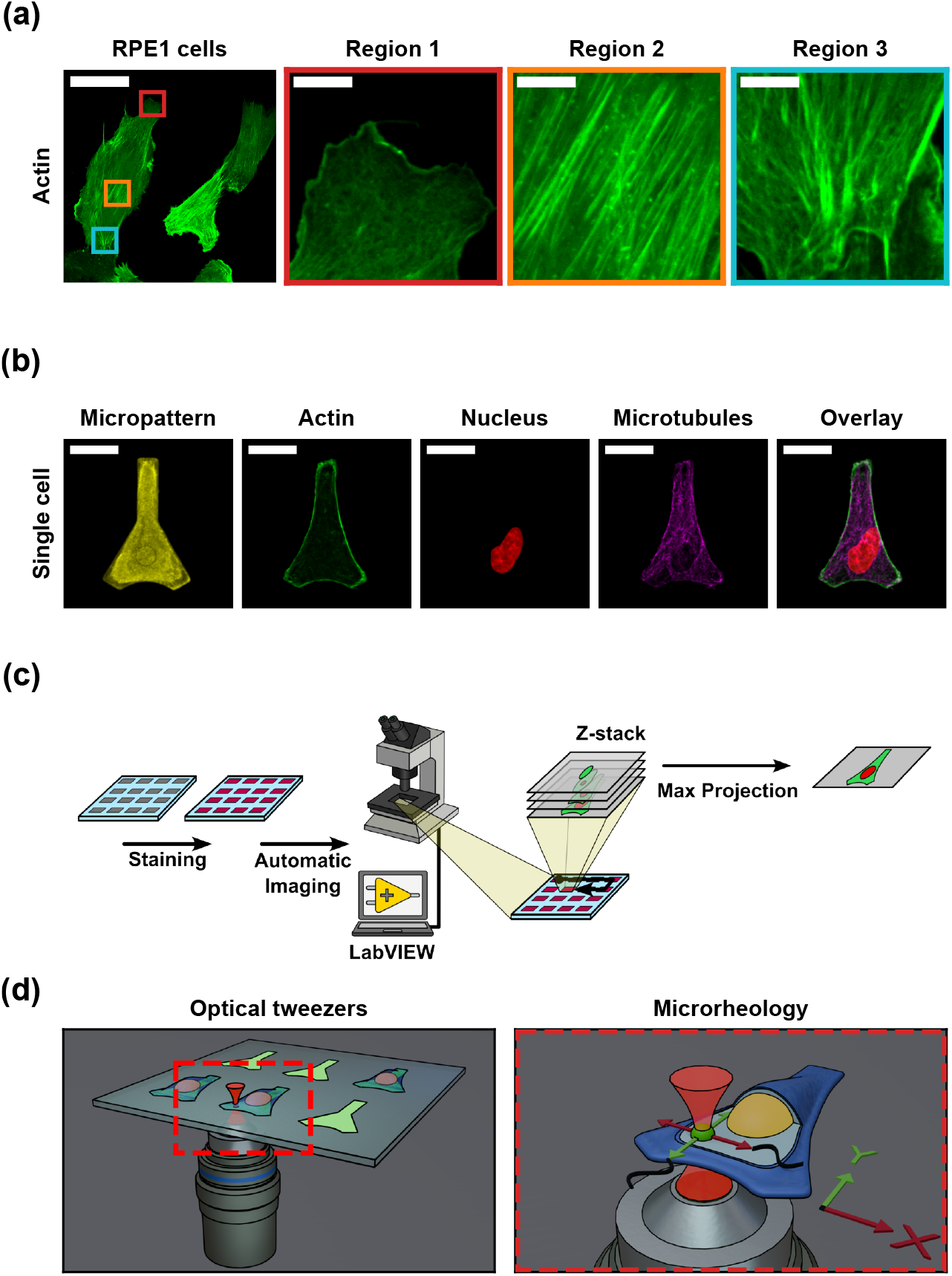
Workflow for mapping intracellular structural and mechanical heterogeneity and anisotropy. **a**: Representative unconfined RPE1 cells with zoomed-in regions highlighting the local structural heterogeneity and anisotropy of the actin network. Actin filaments form branched network in region 1, whereas regions 2 and 3 show prominent, directionally oriented actin stress fibers. Scale bar in left panel: 80 µm, Scale bars in other panels: 10 µm. **b**: Individual and merged fluorescence channels of an RPE1 cell geometrically constrained on the fibronectin micropattern. Scale bars: 20 µm. **c**: Schematic representation of the immunostaining and automated multi-channel Z-stack image collection pipeline utilized to construct the “Average RPE1 cell” structural template. **d**: Schematic of a microrheology experiment performed by optical tweezers in x and y direction.

To address this question, we first had to overcome two major challenges. The first challenge is that single cells show a high degree of shape variability (Fig. 1a), even within the same cell line. This variance results in the lack of a well-defined spatial reference frame to compare mechanical measurements across different cells. To standardize cellular morphology, cells were confined on a symmetrical, flask-shaped fibronectin micropattern with a concave bottom (Fig. 1b). The geometry features a long (26 µm), narrow (9 µm) vertical neck that transitions into a flared, triangular base with an upward-curving basal arc. The restrictive width of the upper neck mechanically blocks the nucleus, forcing it to remain in the wider base, while the concave bottom and sides of the pattern induce high membrane tension to promote the formation of peripheral actin stress fibers along the boundaries [27]. Using this technique, the global morphology of the cells can be systematically controlled.

The second challenge stems from the fact that local mechanical properties might collectively depend on all structures existing within the cytoplasm. Ideally, we would need to image all these components simultaneously during microrheology experiments. However, this is practically impossible due to the limited number of available fluorescence channels. To overcome this limitation, we alternatively sought to determine the average spatial localization of different structures on this specific micropattern, as introduced previously [28]. To achieve this, we prepared distinct samples across a grid of micropatterns, staining a single specific target, namely microtubules, the Golgi apparatus, or the endoplasmic reticulum (ER) in individual samples, while co-staining actin filaments, nucleus, and the micropattern boundaries across all samples. Subsequently, we utilized a home-built LabVIEW program to automatically capture multichannel Z-stacks of every single cell found on the patterns (Fig. 1c). From this collected image dataset, we generated maximum intensity projections of the individual cells and calculated the average intensity distribution for each structural component, by finding the Wasserstein barycenter, as described in detail below. By overlaying these averaged structures, we successfully constructed an “Average RPE1 cell” template.

With this standardized structural framework established, we next deployed optical tweezers to perform active and passive microrheology experiments (see Materials and Methods), quantifying the mechanical fingerprint that represents the complex shear modulus and mechanical activity in both the *x* and *y* directions (Fig. 1d). Because the 1 µm large polystyrene beads used as microprobes are randomly distributed throughout the cytoplasm, aggregating the individual data points from many cells allows us to span the entire geometry and generate a twodimensional mechanical map. By comparing the mechanical map with the average RPE1 cell, we can then systematically explore any potential correlations between these localized mechanical properties and the localization of different intracellular structures.

## 3 Wasserstein Barycenters as Average Cell Template

The most straightforward approach to constructing an average RPE1 cell template is to align the individual images using the micropattern channel and subsequently calculate their pixelwise linear average. However, this conventional method often introduces unphysical artifacts. For instance, in the case of actin filaments, linear averaging applied to a low number of images yields multiple artificial cell boundaries due to minor discrepancies of spatial positioning of the boundary between individual cells (Fig. 2a, left). As the number of images increases, the average image becomes blurred, effectively smearing out finer structural details (Fig. 2a, right).

**Figure 2:**
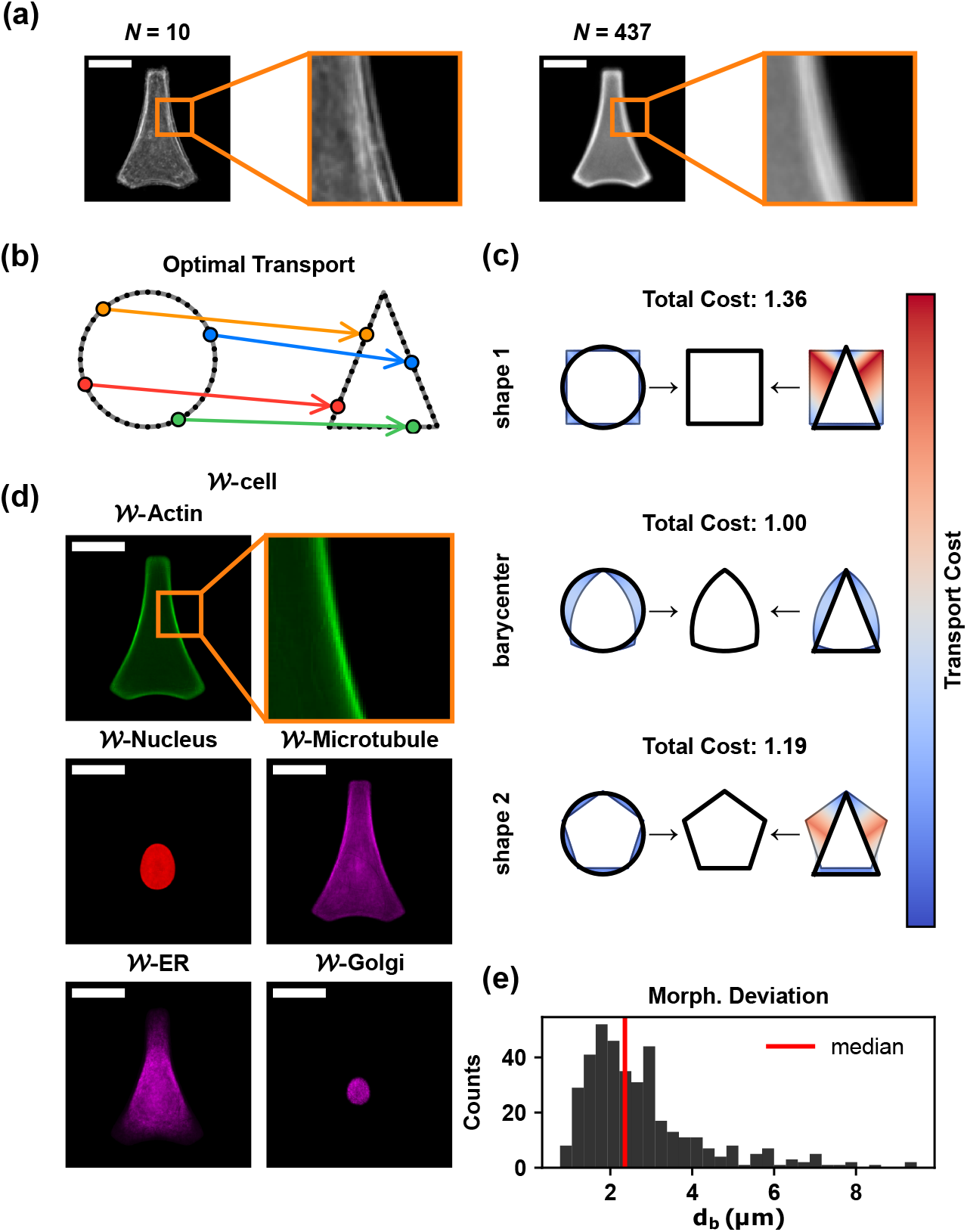
Optimal transport framework to construct average RPE1 cell. **a**: Linear pixel-wise averaging of actin images (N = 10 and N = 437), demonstrating typical artifacts including multiple boundaries and feature blurring. Scale bars: 20 µm. **b**: Schematic representation of optimal transport mapping, illustrating the displacement required to deform a source shape (circle) into a target shape (triangle). Arrows show how four representative points should be transported optimally. **c**: The Wasserstein barycenter represents the optimal shape that minimizes the cumulative cost of transporting individual shapes (here, a circle and a triangle) into a shared average representation. Three consecutive rows compare the barycenter against two alternative candidate midpoint shapes, a square and a pentagon. The transport pathways connecting each individual shape to the midpoint geometries are color-coded by their transformation costs, normalized to the total cost of the barycenter. This visualization demonstrates that the barycenter achieves the minimum total transformation cost among all possible geometric configurations. **d**: Calculated barycenters of different structures, including actin, nucleus, microtubules, the Golgi apparatus and ER. Scale bars: 20 µm. **e**: Histogram of the Wasserstein distances of individual actin images relative to the actin barycenter (d_b_), quantifying the divergence of single cells from the population median.

To avoid these artifacts we require a different notion of interpolating and averaging between images that takes into account deformations of the cell structures. A natural choice is the Wasserstein distance, as induced by optimal transport [29]. In this framework one interprets images as probability densities and asks for the most efficient way to transport one into the other. This provides a mathematically sound notion of distance and continuous interpolation between two given probability distributions and also a way to compute the average distribution of a whole dataset.

For a concrete illustration, consider two different shapes, like a circle and a triangle, each discretized by an equal finite amount of points (Fig. 2b). To obtain the optimal transport distance between them one must consider all possible assignments between the two point clouds (here corresponding to permutations) and identify the one with the lowest cost. Here, the cost of deformation is defined as the average squared distance by which all points are moved. In Fig. 2b, this minimum-cost assignment is visualized with arrows that indicate how points move from initial to final configuration.

The average distribution, known as the Wasserstein barycenter, is then defined as the distribution for which the total optimal cost of deformation from the barycenter to all individual distributions in the dataset is minimized. In Fig. 2c, the barycenter is compared to two other possible average shape candidates, a square and a pentagon. The pathway for transporting each point to its destination, is color-coded by the squared distance between the points. As shown, the paths for the true barycenter remain blue, leading to a lower total cost compared to the other candidate shapes.

By applying this method to our set of cell images for each individual structure, we can compute a barycenter that serves as the average structural representation, hereafter denoted by the prefix *W*-(e.g., *W* -actin and *W* -nucleus, with *W*-cell representing the combined overlay of these *W*-structures). This approach preserves characteristic features better than conventional linear averaging. As shown in Fig. 2d, the *W*-actin displays a single sharp boundary that truly represents the actin architecture observed in single cells (Fig. 1b). Moreover, the *W*-nucleus possesses a sharp border compared to the blurred linear average (SI Fig. 1), while reflecting the geometric constraints of the micropattern by appearing horizontally centered and vertically symmetric and elongated. The *W*-microtubule is concentrated near the nucleus, reflecting the location of the microtubule-organizing center (MTOC), while also colocalizing with actin stress fibers on cell periphery and uniformly filling other regions of the cytoplasm. Furthermore, - ER indicates that this organelle occupies a substantial portion of the total cell volume, with its highest density concentrated in the perinuclear region. This closely mirrors the spatial localization of the ER observed within individual cells (SI Fig. 1). Consequently, this finding highlights the potential contribution of the ER in global mechanical properties of the cytoplasm [30–32], which remains mostly overlooked in most previous studies.

There is however one notable caveat to this *W*-ER as it artificially bleeds into the nuclear region, whereas individual cells exhibit a signal void where the nucleus resides (SI Fig. 1). A similar but more severe discrepancy occurs with the average *W* -Golgi, which fails to capture the perinuclear localization of this organelle observed in individual cells (SI Fig. 1). This inaccuracy arises because the barycenter of each structure is calculated independently. Therefore, for organelles such as the Golgi apparatus and ER which are coupled to the nucleus, a joint probability distribution with the nucleus might be required. However, this discrepancy also occurs when using the conventional linear averaging, which similarly suffers from disconnection between the average representation and the true localization of the Golgi in individual cells, while simultaneously blurring the entire average image (SI Fig. 1).

It is important to note that although the barycenter can serve as average template for the cellular structures, individual cells exhibit substantial deviations from this average configuration. This cell-to-cell variability of cellular shape is quantified by the distribution of the Wasserstein distances between the individual single-cell instances and the barycenter (*d*_*b*_) calculated for actin (Fig. 2e) and other structures (SI Fig. 2).

## 4 Heterogeneity and Anisotropy of Mechanical Properties and Activity

To characterize the mechanical properties of the RPE1 cytoplasm, we performed microrheology experiments (SI Fig. 3) on 277 microprobes across 133 cells. Because the microprobes were randomly distributed within the cytoplasm of the individual cells, the aggregated dataset structurally spanned the majority of the cytoplasmic domain (Fig. 3a). To avoid spatial misalignment caused by cell-to-cell variability, we transformed individual probe coordinates into the standardized *W* -cell space using Gaussian process interpolation (see Materials and Methods). This means the position of the particles was corrected using the Wasserstein barycenter-based vector fields. Additionally, we mirrored the data points across the symmetry axes to reflect the inherent geometric symmetry of both the micropattern and the *W* -cell template. It is important to note that any microprobe coordinates mapped within the projected nuclear region does not represent intranuclear mechanical properties; rather, they correspond to cytoplasmic microprobes physically positioned above the nucleus along the *z* dimension. These microrheology experiments yield the complex shear modulus *G*^*∗*^(*f*) and the effective energy spectrum *E*_eff_(*f*) (Fig. 3b-c). The modulus *G*^*∗*^(*f*) is a complex-valued quantity, whose real part, the storage modulus *G*^*′*^(*f*), quantifies the elastic behavior of the material, whereas its imaginary part, the loss modulus *G*^*′′*^(*f*), reflects the viscous properties.

**Figure 3:**
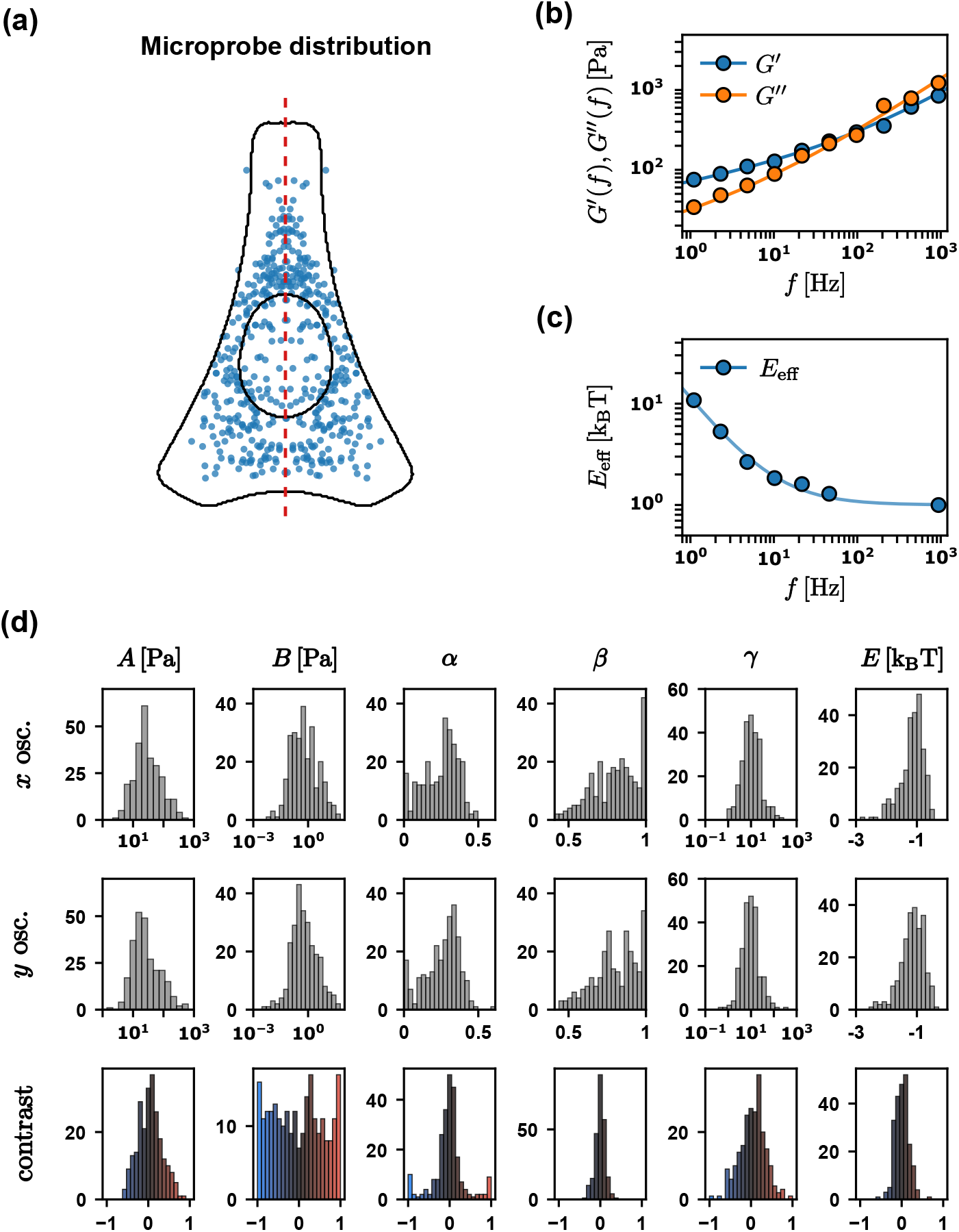
Microrheology and mechanical heterogeneity. **a**: Distribution of measured microprobes within the cytoplasm. Due to the symmetry of the pattern, the probe position is also mirrored along the pattern’s symmetry. The border of W -actin and W -nucleus is also shown (N_cells_ = 133, N_particles_ = 277). **b**: A representative storage modulus G^′^(f) and loss modulus G^′′^(f), as measured by active microrheology. The solid lines indicate the fitted fractional Kelvin-Voigt model. **c**: A representative effective energy spectrum estimated by comparing the dissipative response χ^′′^(f) measured by active microrheology with the one estimated by FDT. The solid line indicates the fitted power-law model. Values in the range from 50 to 500 Hz are omitted as here the noise due to the spinning disk dominates the free particle motion. **d**: Statistical distributions of the viscoelastic fit parameters (A, B, α, β, E, and γ) measured along the x-direction (first row, n_x_ = 260) and y-direction (second row, n_y_ = 257). The third row displays the parameter contrast between the two orthogonal directions (n_contrast_ = 240). Note that the total number of accepted measurements along the x and y axes is lower than the total number of measured particles, as cases yielding high noise in the calculated G^∗^(f) or the effective energy spectra E_eff_(f) were excluded. To visualize potential directional contrast, a color map is used which scales continuously from blue to black to red, corresponding to a parameter contrast range of −1 to 0 to 1.

To parametrize these mechanical properties, we fitted each measured *G*^*∗*^(*f*) curve (solid lines in Fig. 3b) using a fractional viscoelastic model [33, 34]

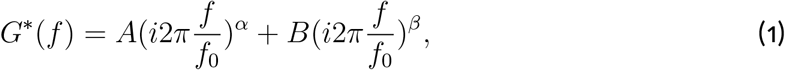

where *f* represents the driving frequency and *f*_0_ = 1 Hz. This model yields four distinct fitting parameters (*A, B, α*, and *β*), where *A* and *B* quantify the solid-like and fluid-like scale factors (or mechanical resistance in general), while the power-law exponents *α* and *β* indicate the extent to which the viscoelastic material behaves like a solid versus a fluid. Specifically, an exponent of 0 denotes a purely elastic solid, whereas an exponent of 1 represents a purely viscous fluid [11]. Incorporating two distinct power-law terms is necessary to capture the crossover feature between *G*^*′*^(*f*) and *G*^*′′*^(*f*) (Fig. 3b), which is a behavior frequently observed when probing the complex mechanics of the intracellular environment [20, 35, 36]. Moreover, the effective energy also behaves like a power-law (Fig. 3c). We therefore fit a second power-law function

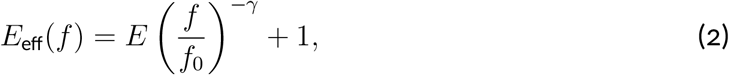

which yields two further parameters *E* and *γ*. The scale factor *E* quantifies the activity of the environment surrounding the microprobe, while the exponent *γ* indicates how rapidly this activity decays at higher frequencies. Together with the four parameters described before, they form an active mechanical fingerprint [20].

Finally, to quantify the anisotropy of this mechanical fingerprint, measurements are performed along both the *x* and *y* axes for each microprobe. Then to quantify anisotropy, we define the contrast for mechanical parameters as follows

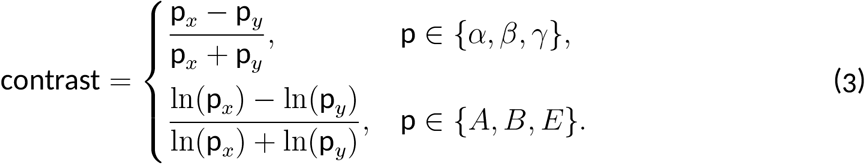

We calculate the contrast only for particles where measurements in both *x* and *y* directions are available. The statistical distributions of all measured rheological parameters and their respective spatial contrasts are presented in Fig. 3d. The scale factors *A, B*, and the effective energy *E* exhibit log-normal distributions, whereas the power-law exponents *α, β*, and *γ* follow rather a normal distribution. Intriguingly, the log-normally distributed parameters display extreme heterogeneity, varying across at least two orders of magnitude. The normally distributed parameters likewise span a remarkably broad portion of their physically permissible domains. Specifically, the exponents *α* and *β* cover nearly the entire lower and upper halves of the [0, 1] interval, respectively.

To evaluate potential global directional bias, we performed a statistical *t*-test on each parameter dataset comparing the *x*- and *y*-directions. None of the parameters exhibited a statistically significant difference, with *p*-values all exceeding 0.25. This global ensemble symmetry is further reflected in the average parameter values along the orthogonal axes, with *Ā*_*x*_ = 50 Pa closely matching *Ā*_*y*_ = 48 Pa, and the effective energy tracking symmetrically at *Ē*_*x*_ = 16.9 k_B_T and *Ē*_*y*_ = 15.7 k_B_T.

However, analyzing the parameter contrast on a per-microprobe basis reveals striking structural differences. While the power-law exponents show zero contrast, tightly accumulating around 0, the scale factors *A, B*, and the energy *E* display highly heterogeneous contrast profiles. To ensure these variations reflect physically meaningful anisotropy rather than measurement artifacts, we quantified the intrinsic directional bias of our setup by performing microrheology on an isotropic viscoelastic hydrogel (SI Fig. 4). These control experiments revealed a slight bias of 20 % toward higher values of parameter *A* along the *y*-axis, corresponding to a baseline instrument contrast of

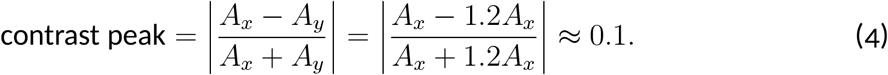

Because the observed contrast distribution tails extend far beyond this ∼0.1 instrumental offset, the recorded contrast reflect true cellular anisotropy rather than experimental artifacts. Based on these statistical distributions, we suggest that the cytoplasm is characterized by an intriguing dual nature, i.e. while it exhibits global isotropy at the ensemble level across cells, it is strictly governed by extreme heterogeneity and sharp anisotropy at the local level. By evaluating these parameters together as an integrated active mechanical fingerprint, we suggest the following physical and biological interpretations of our data.

First, by convention, *α* is constrained to be the smaller exponent (*α < β*). Consequently, the first power-law term in Eq. 1 dominates the material response at lower driving frequencies, whereas the second term becomes dominant at higher frequencies. Given that the ensemble averages and particle-wise contrasts reveal no directional anisotropy in either exponent, we argue that the cytoplasm’s predominant physical behavior is best described as solid-like at low frequencies and transitions to a fluid-like medium at high frequencies, independent of the direction probed. However, the exact extent to which any given particles feels a solid-like or a fluid-like environment, as defined by the actual values of the power-law exponents in a particular cell, remains highly heterogeneous, which is reflected in the broad distributions of *α* and *β*. This highlights that while the cytoplasm as seen globally across many cells, averages out to an isotropic continuum on an ensemble level even despite the reproducible cell shape. In contrast, the specific viscoelasticity experienced by an individual organelle or macromolecular complex in a single cell remains hyper-variable and highly unpredictable due to this local heterogeneity.

However, the picture changes when focusing on the prefactors *A* and *B*. Here we interpret the contrast profile of these two (Fig. 3d) as direct evidence that on the local level of the probe particles in a single cell, the mechanical properties of these individual micro-domains are highly anisotropic. This can be seen in the large contrast measurements that cannot be explained by any anisotropy originating from the instrument. Such anisotropy suggests that when a microprobe is embedded in the cytoplasm, its immediate surroundings can be anisotropic, while this anisotropy averages out across many cells, and is hence not an outcome of the cell shapes. Therefore, the probe experiences an environment where the stiffness or viscosity can be higher in one direction than the other. Because these directional variations emerge randomly at local level, they blend into a global isotropy when pooling the measurements together at the ensemble level. However, at the individual probe level, the sharp directional anisotropy remains completely preserved.

Surprisingly, the wide contrast distribution of *E* implies that while active metabolic forces vary drastically, the resulting active fluctuations show a random anisotropy. A possible explanation for this local anisotropy within a given cell is that the direction-dependent stiffness reflects the intracellular organization at the specific measurement location. Thus, our measurements may probe intracellular organization from a mechanical perspective, even though this local organization does not necessarily recapitulate the overall cell architecture imposed by the micropattern. In contrast, the overall physical properties remain surprisingly isotropic in global level.

## 5 Major Descriptors of Intracellular Mechanics

To determine which of the six parameters capture the primary variation in intracellular mechanics, we evaluated the Pearson correlation coefficient for all parameter pairs independently for the *x* and *y* directions (SI Fig. 5). The analysis revealed that the parameters are not fully independent. In particular, parameters *B* and *α* exhibit strong correlations, while *B* and *β* are anti-correlated, confirming previous findings that had been obtained across different cell types [20]. To account for these dependencies, we first standardized each parameter p by computing its *z*-score *z*_*i*, p_

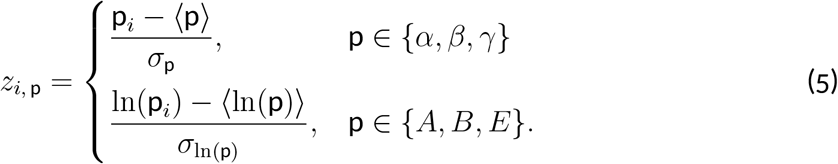

Here, *i* indexes the individual value of parameter p, ⟨·⟩denotes the population mean, and *σ* represents the standard deviation. The resulting *z*-score quantifies the deviation of observation *i* from the population mean in units of standard deviation. We then performed principal component analysis on the standardized parameters. The first three principal components, *C*_1_ −*C*_3_ account for nearly 85 % of the overall data variability (SI Fig. 6a, b, left panels). Analyzing the component contributions shows that *C*_1_ is heavily dominated by *B, α*, and *β*; *C*_2_ is governed by *E* and *γ*; and *C*_3_ is primarily captured by *γ* (SI Fig. 6a, b, right panels). Because *B* and *β* are anti-correlated while *β* and *α* are positively correlated, an increase in *C*_1_ corresponds to a transition toward more fluid-like material behavior. Consequently, following reference [20], we define *C*_1_ as the fluidity component. Moreover, although *C*_2_ captures the variation of active forces through both *E* and *γ*, we focus only on *E* as has been done previously [20] to represent the active forces, which isolates overall activity. Unlike in prior work [20], where *C*_3_ was captured primarily by *A, γ* displays higher variability in patterned cells and strongly affects *C*_3_. Nevertheless, since mechanical activity is already accounted for by *E*, we maintain the established parameter set (*A, E, C*_1_). Replacing *A* with *γ* would omit mechanical resistance from our phase-space, whereas combining mechanical resistance (*A*), activity (*E*), and fluidity (*C*_1_) defines a balanced phase-space.

## 6 Spatial Mapping of Intracellular Mechanics and Activity

Having characterized the global ensemble distributions and probe-wise contrasts of the mechanical fingerprint, a compelling question arises: Can this pronounced heterogeneity and anisotropy of the mechanical fingerprint be mapped onto distinct spatial regions of the cytoplasm? Given that the organelles and cytoskeletal networks exhibit well-defined, non-uniform localization patterns within the *W*-cell template (Fig. 2d), one might intuitively hypothesize that specific regions, such as the densely packed perinuclear zone or the highly aligned peripheral margins, would systematically display different directional mechanical properties.

To test this hypothesis and systematically search for spatial correlations, we interpolated our spatially discrete measurements into a continuous map across the cytoplasmic domain using two-dimensional smoothing splines (Fig. 4). The spatial maps for these parameters appear remarkably uniform and flat. When processing our datasets using radial basis function interpolation to generate continuous contour maps, we indeed observe distinct spatial patterns across the cellular domain (SI Fig. 7). However, such smooth interpolation schemes inherently force continuous patterns onto discrete, highly scattered microprobe data regardless of whether underlying spatial gradients truly exist.

**Figure 4:**
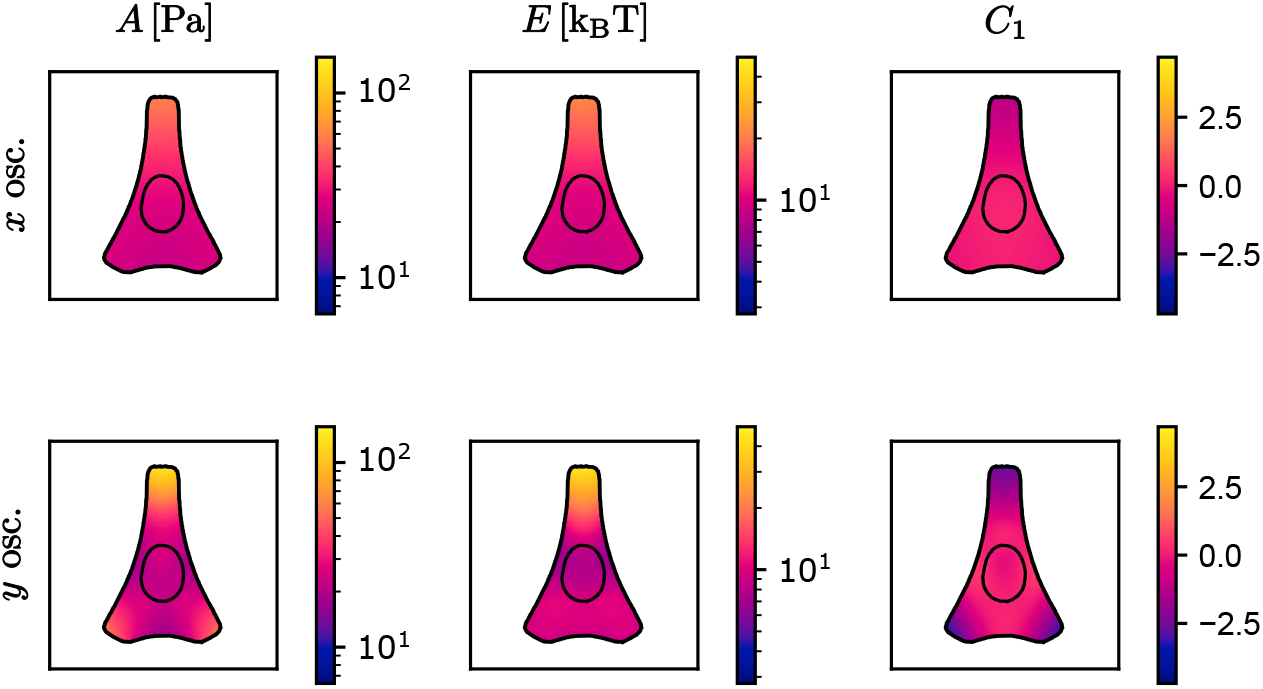
Spatial mapping of mechanical properties and activity across the cytoplasm. Continuous twodimensional smoothing splines fitted to the spatial distributions of A, E, and C_1_. The first and second rows present the independent spline fits for measurements performed along the x- and y-axes, respectively. Scale bars: 20 µm.

To test whether spatial features from fitting splines to our microprobe data reflect physical spatial structure, we use a model comparison procedure (see Materials and Methods). We compare how well spline models and spatially invariant models predict data points that were left out of the fit (out-of-sample predictive power, assessed by cross-validation): If a spline model systematically improves these predictions, the spatial structure it encodes provides evidence for physically distinct regions within the cell. If, however, it cannot outperform a constant model containing no spatial dependence, the data provide no evidence that the fitted spatial structure is more than an artifact of measurement noise. To assess this predictive power, we performed nested cross-validation, iteratively separating training data from entirely unseen test data. The spline and constant models compete only on their ability to predict observations not used during either fitting or model tuning, and we compare their performance by the resulting out-of-sample prediction error.

Across the nested cross-validation iterations, the spline models show no significant improvement in out-of-sample predictive performance over the constant baseline. Our data therefore provide no evidence for reproducible spatial gradients or regional localization at the scale resolved by these measurements. In particular, we cannot resolve systematic differences in the mechanical scale factors or power-law exponents between microprobes positioned near the cell periphery and those embedded within the perinuclear region.

This means that a microprobe positioned near the cell periphery detects, on average, the same mechanical scale factors and power-law exponents as one embedded within the perinuclear region. Furthermore, the variance and data spread observed in our global histograms remain nearly uniform across the entire cellular geometry (Fig. 4). This spatial invariance demonstrates that the global localization of organelles and cytoskeletal architectures seen in the *W*-cell template does not automatically translate into global distinct mechanical zones measured by a 1 µm microprobe size. This suggests that either the cells operate on a flat global mechanical landscape, providing organelles with a predictable mechanical baseline regardless of location, or that global mechanical gradients do exist but are too small to be distinguished from the local mechanical variability in our measurements, as discussed below.

## 7 Origins of Spatial Uniformity: Micro-scale Structures and Cell-to-cell Variance

The lack of distinct spatial gradients in our mechanical maps presents an interesting paradox: how can a cytoplasm with such visually distinct structural regions maintain spatially uniform average mechanical map? We argue that this spatial uniformity does not necessarily imply that the cytoplasm is an intrinsically uniform material on the global level. Rather, this uniform spatial map potentially arises from local structural heterogeneity and anisotropy captured by our 1 µm microprobes.

To evaluate this paradox, we first determined whether our experimental setup possessed sufficient spatial resolution to capture potential global mechanical gradients. To this aim, we computed the spatial correlation coefficients from individual fluorescent images of actin, microtubules, the Golgi apparatus, and the endoplasmic reticulum (Fig. 5a). Each cellular structure exhibited a distinct characteristic length scale that is a measure of at which length we expect significant structural variations of each intracellular component; when combined, the ensemble average revealed a structural correlation length of *l*_*c*_ = 3.84 µm at the 1*/e* decay threshold. This correlation length represents an upper limit for the microprobe diameter, which means that probes significantly larger than *l*_*c*_ would average out global mechanical features and artificially flatten spatial gradients. Because our microprobes are 1 µm in diameter, which is substantially smaller than *l*_*c*_, our measurements retain sufficient spatial resolution to detect the here reported large intracellular mechanical variations.

**Figure 5:**
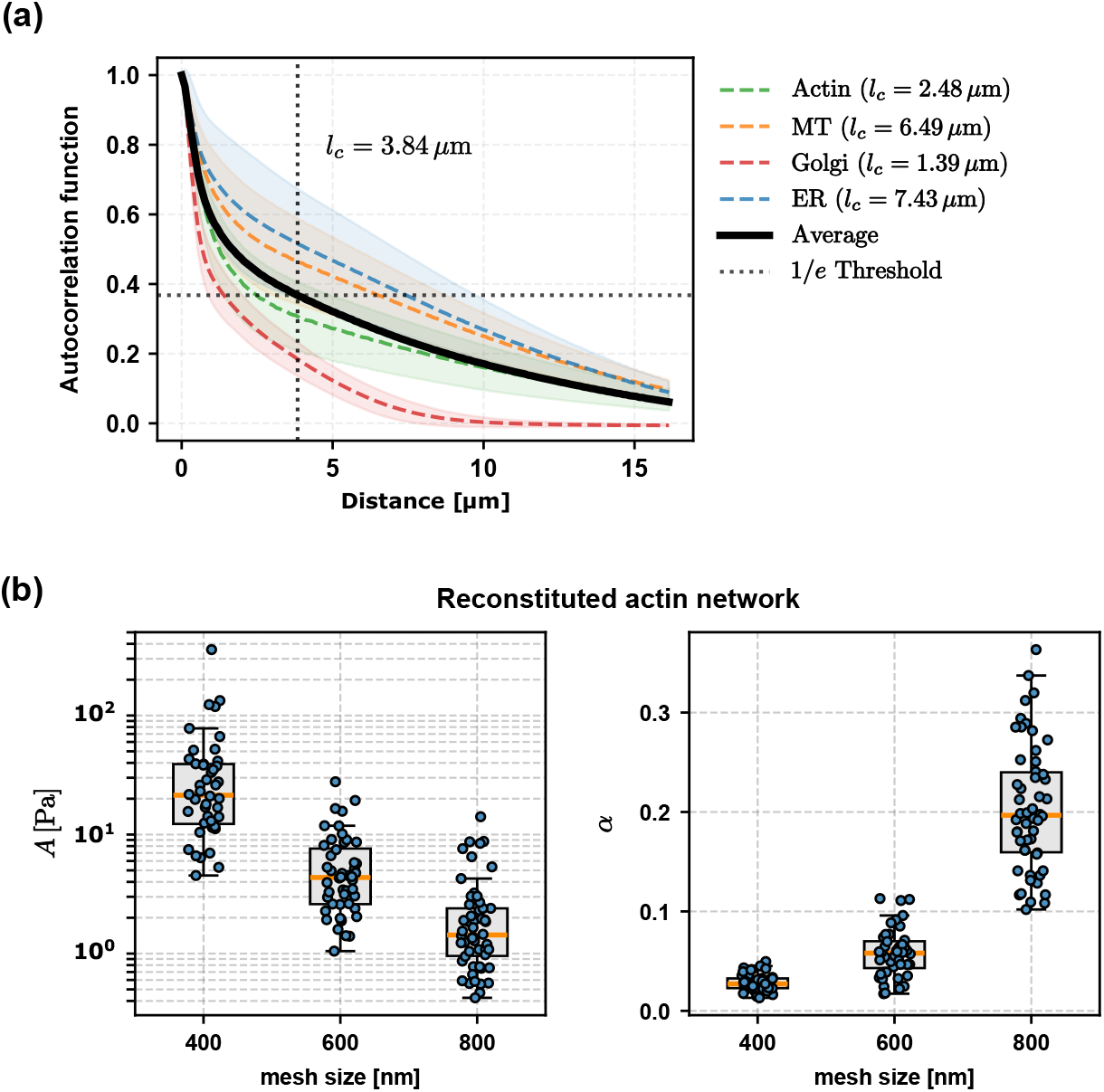
Verifying spatial resolution of the instrument. **a**: Spatial correlation coefficients l_c_ calculated from individual images of distinct cytoplasmic structures as a function of distance. The solid black curve represents the global mean of the individual structural correlation profiles, reaching a characteristic 1/e decay threshold at 3.84 µm. This length scale defines the maximum microprobe dimension required to capture an effective, averaged structural continuum. **b**: Parameters A and α measured in reconstituted actin networks with varying mesh size.

To confirm that our probes can resolve spatial variations in mechanical properties, we performed microrheology experiments in reconstituted actin networks with varying mesh sizes *ξ*. We chose here deliberately actin networks as test structure, as we know from previous experiments [20] that actin networks are not the dominating mechanical structure for intracellular mechanics. In turn it is a standard biopolymer system with viscoelastic properties comparable to the observed intracellular situations. As we test here the possibility of our measurement to reliably detect the spatial variability, such biopolymers are good reconstituted control environments. Because the microprobe diameter remained constant across all conditions, varying *ξ* simulates spatial variations in network density. As shown in Fig. 5b, networks with different mesh sizes exhibit distinct mechanical behaviors. In fact, as the mesh size increases, parameter *A* decreases significantly, whereas parameter *α* increases. This increase in *α* indicates a transition from a solid-like to a fluid-like regime. All changes across adjacent mesh size pairs (400 nm–600 nm and 600 nm–800 nm) were statistically significant (*p <* 0.001, Student’s *t*-test).

Having it verified that we have enough spatial resolution and we can capture mechanical differences, we should answer the question why no meaningful gradient of mechanical properties is observed. In fact, any potential global mechanical gradients may be masked by the heterogeneity driven by cell-to-cell variability. To investigate whether cell-to-cell variability is present in our dataset, we grouped the data points by individual cell instances and sorted the cells based on their median *A*_*y*_ values (Fig. 6a). While a clear trend emerged across sorted cell medians, our experiments do not allow calculating a reliable average of mechanical properties for each individual cell due to the fact that most cells contained only one or two microprobes. However, in the few instances where cells contain at least four microprobes, their median stiffness values might differ by up to an order of magnitude (two of such instances are highlighted by green and yellow colors in Fig. 6a.

**Figure 6:**
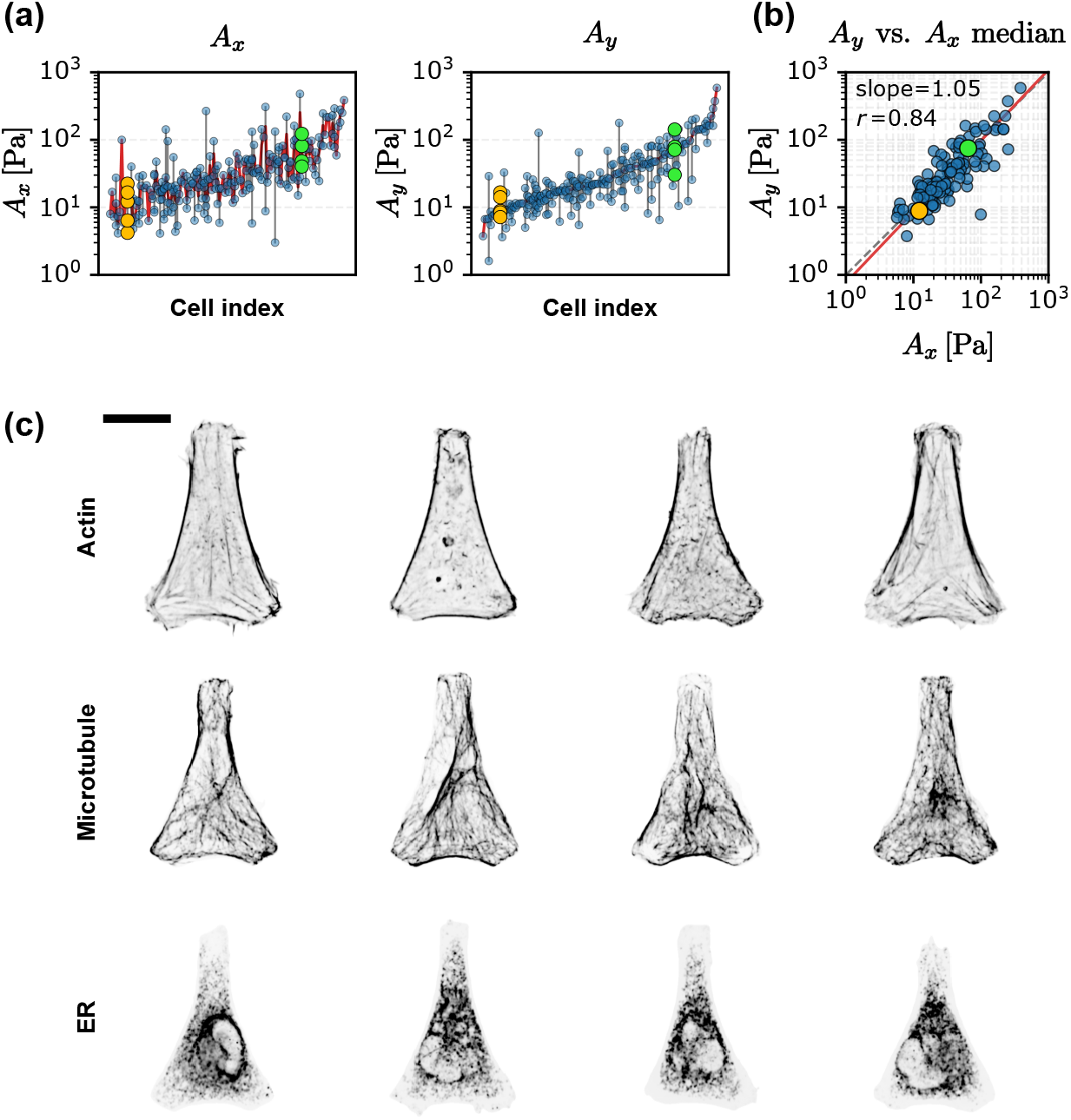
Cell-to-cell variability. **b**: Analysis of cell-to-cell mechanical variability. Microprobe measurements of the scale factors A_x_ and A_y_ are grouped per individual cell. The cellular datasets are ordered along the horizontal axis based on the sorted median value of A_y_ for each cell (red trend line), with the corresponding A_x_ values plotted under the identical cell sorting (blue data points). The distinct trend in A_y_ highlights a drastic cell-to-cell variability (data points of two representative cells are shown in green and yellow), while the higher fluctuations of A_x_ is a result of profound anisotropy in each cell. **b**: Scatter plot comparing the median A_y_ versus median A_x_ values calculated for individual cell instances. The solid line represents a major axis regression fit yielding a slope of 1.05. The data spread perpendicular to the regression line highlights mechanical anisotropy within individual cells. **c**: Representative fluorescence images demonstrating cell-to-cell structural variability across a grid of three distinct components (rows: actin filaments, microtubules, and endoplasmic reticulum) and four individual cell instances (columns). Pronounced intracellular architectural reorganization is visible between instances despite the preservation of global cellular morphology enforced by the micropattern. Scale bar: 20 µm.

We then wondered if a cell showing a systematically higher stiffness in *y* axis provide an overall higher stiffness offset. To answer this, we checked if the stiffness as presented by the parameter *A* scales similarly along the *x* and *y* axes. As shown in Fig. 6a, if we sort *A*_*x*_ values using the sorted indices from *A*_*y*_, we see a similar trend but with more noise coming from local anisotropies. A scatter plot comparing the cell-median values of *A*_*x*_ and *A*_*y*_ (Fig. 6b) shows a clear correlation with a regression function that provides an almost perfect linear scaling of 1.05. This scaling indicates that a cell that is stiffer along one orthogonal axis is proportionally stiffer along the other. Nevertheless, the spread perpendicular to the regression line confirms that significant local anisotropy persists within individual cells.

Together, these findings point to cell-to-cell variability as a potential driver of data variability, arising primarily from both structural fluctuations and cell-cycle variations. First, stiffness scales as a power law of mesh size in biopolymer networks made up of single components such as Factin

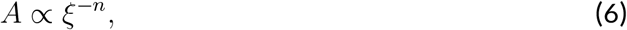

where *n* varies from 3 for networks of semi-flexible chains to 5 for cross-linked networks. [37]. Our reconstituted network data agrees well with *n* = 4, where average stiffness *A* drops from 39.1 to 2.4 Pa (Fig. 5b). Differentiating Eq. 6 yields the variability of *A* to local mesh size fluctuations

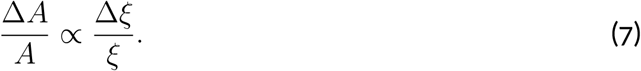

This relation only provides a controlled framework to understand how structural fluctuations translate into data variability, demonstrating that greater mesh size variability directly results in increased spread of stiffness. In contrast, it does not isolate or dissect the specific role of actin network in intracellular mechanics. When turning to the cells, they display substantial structural variations even when confined to identical micropatterns (Fig. 6c), which is also reflected in the broad distribution of Wasserstein distances (Fig. 2e). Therefore, the local cytoplasmic structure and composition vary significantly across cell population at any given position relative to the micropattern, which leads to the pronounced error bars in the spatial correlation functions (Fig. 5a). Consequently, we likewise expect the measured mechanical stiffness across the cell population to exhibit broad spatial fluctuations. Second, because our experimental cell populations were non-synchronized, temporal physiological differences introduce an additional, independent layer of data variability. Because, beyond the mechanical changes known to occur during mitosis [36], cells undergo continuous mechanical modulation across sub-stages of interphase [38].

In summary, while our microprobes provide sufficient spatial resolution to detect intracellular mechanical gradients, these spatial variations might be effectively masked by broad cell-to-cell variability driven by structural heterogeneity and cell-cycle non-synchronization.

## 8 Conclusion

In this work, we developed a structural and mechanical mapping framework that combines linearized optimal transport with active and passive optical tweezers microrheology, to investigate intracellular mechanical heterogeneity and anisotropy. We showed that RPE1 cells exhibit highly organized, non-uniform average spatial positioning of organelles and cytoskeletal networks when constrained to an adhesive micropattern. Conversely, although the measured mechanical properties and activities are locally highly heterogeneous, spanning over more than two orders of magnitude, the corresponding spatial maps of the averages within the cells appear uniform within the large statistical variability.

We argue that this apparent uniformity and data dispersion are driven by cell-to-cell variability. By calculating the spatial autocorrelation coefficient across different cellular structures and characterizing the mechanical properties of reconstituted actin networks with varying mesh sizes, we confirmed that our experimental setup possesses sufficient spatial resolution to detect mechanical gradients. However, cell-to-cell structural variability introduces a high degree of spread, consistent with other studies where a high cell-to-cell variability of mechanical properties has been reported [15]. This variability might effectively mask any underlying global mechanical gradients. Consequently, ∼1 µm intracellular objects and organelles experience a highly stochastic local environment driven by cytoplasmic heterogeneity and anisotropy. Furthermore, variations in cell cycle stage among non-synchronized cells likely contribute to these mechanical fluctuations, driving both structural and mechanical variability.

Future studies could isolate individual contributors to cell-to-cell variability for instance, by arresting cells in a specific cell-cycle stage, such as G1 [39]. Combining synchronization with emerging techniques like mean back relaxation [40] and smaller microprobes would enable a statistically robust number of measurements, facilitating spatial mechanical mapping for individual cells. Furthermore, tracking cells over several hours would capture the temporal evolution of these maps, while high probe number per cell would allow average mechanical properties to be compared across the population to assess cell-specific variations. Ultimately, this approach would enable spatial variations to be systematically decoupled from temporal and cellspecific fluctuations. Importantly, because the microprobes were internalized via endocytosis, they remained encapsulated within a lipid membrane, providing a physical model mimicking native membrane-bound organelles of comparable size. Moving forward, surface modification of these probes to target specific subcellular structures or cytoskeletal networks would enable precise probing of how distinct intracellular architectures drive local mechanical heterogeneity and anisotropy.

## Author Contributions

M.A.E., B.S., and T.B. conceived and designed the research, M.A.E. and D.C.Z. performed active and passive microrheology experiments, D.M. provided the microrheology data analysis pipeline and the viscoelastic hydrogel, S.L. performed statistical modeling and wrote the corresponding Materials and Methods sections, H.S. and M.A.E. performed cell culture, micropatterning, and micropatterning protocol optimization, S.U. computed the *W*-cell barycenters from images captured and preprocessed by M.A.E, B.S. provided the libraries for optimal transport calculations, S.K. provided ideas on how to perform actin measurements. C.M. passivated microprobes for reconstituted actin network measurements, S.D. prepared the actin networks, R.H. provided the GPU-accelerated deconvolution code, B.V. contributed to developing the Lab-VIEW control program for microrheology, M.A.E. developed the automated LabVIEW program for multi-channel image acquisition, M.L. contributed core analytical ideas for data interpretation, M.A.E. and T.B. wrote the manuscript with input and revisions from all authors.

## Acknowledgments

This work was funded by the DFG Project-ID 449750155 – RTG 2756, Projects B3 (to T.B. and M.A.E), A2 (to S.K., S.D. and C.M.) and A7 (to S.K. and R.H.). D.C.Z. has received financial support from the Secretaría de Ciencia, Humanidades, Tecnología e Innovación (SECIHTI). The instruments used by M.A.E., D.C.Z and T.B. were funded by the DFG, project IDs 456112451 and 492390964. B.E.V. and T.B. have received funding from the European Research Council (ERC) under the European Union’s Horizon 2020 research and innovation programme (PolarizeMe, grant agreement no. 771201) and by the Deutsche Forschungsgemeinschaft (DFG) under Germany’s Excellence Strategy (EXC 2067/1-390729940). T.B. has received funding from DFG grant 516046415. S.U and B.S. are supported by the Emmy Noether Programme of the DFG, project ID 403056140. Also, D.M, and T.B. have received funding from the DFG, project ID 516046415. We thank Kerstin von Roden for help with establishing the micropatterning protocol, and Ulrike Schulz for preparing the cell line and engineering it to express fluorescence using a lentiviral technique.

## Declaration of Generative AI and AI-assisted Technologies

M.A.E. used Gemini in order to improve the language, flow, and clarity of the manuscript. S.L. used Claude Code (Anthropic) in order to refactor code and fix bugs. S.L. used Grammarly and Claude (Anthropic) to perform spell-checking and improve the manuscript’s language. After using these tools, the authors reviewed and edited the content as needed and take full responsibility for the content of the published article.

## Data Availability Statement

Experimental data will be available from:

https://data.goettingen-research-online.de/

Analysis code will be available from:

https://github.com/m-a-eskandari/avg_cell_mech_mapping/

## Materials and Methods

### Micropatterning

Micropatterns were created following a previously published method [41]. The process includes four main steps

#### Cleaning

18 mm ×18 mm no. 1.5 coverslips were used as the substrate. They were thoroughly washed by submerging them in a dish filled with 99 % ethanol and sonicating them for 15 min. The coverslips were then dried with pressurized air and exposed to plasma for 5 min.

#### Passivation

The cleaned coverslips were passivated immediately after cleaning, using a passivating solution of PLL(20)-g[3.5]-PEG(2) (susos, Switzerland) diluted in 7.4 mM HEPES buffer. To this aim, two spacers made of Polydimethylsiloxane (PDMS) with thickness of ∼0.5 mm were placed parallel to each other on a piece of Parafilm. The coverslips were placed on these two spacers with their main side facing downward. Then, 150 µL of the passivating solution was injected into the space between the coverslips and the Parafilm, and incubated in room temperature for 1 hour.

#### Micropatterning

The coverslips were washed with Milli-Q water and dried by pressurized air. Then 2.5 µL of Milli-Q water per coverslip was added on a chrome photomask (Deltamask, Netherlands). The coverslips were then placed on water droplets so that the passivated side was in contact with chrome side of the photomask. Afterwards, the coverslips were illuminated through the photomask by 250 nm UV light with a power of 400 µW cm^*−*2^. They were washed again with Milli-Q water, dried with pressurized air and kept in a closed petridish for a maximum of 1 week.

#### Protein coating

On the day of the experiment, the passivated coverslips were placed on two parallel PDMS spacers, similar to the passivation step. Then, 150 µL of a bovine fibronectin solution (Sigma Aldrich) diluted in Phosphate-Buffered Saline (PBS) to a final concentration of 50 µg mL^*−*1^ was injected between the coverslips and the Parafilm, and incubated in room temperature for 1 hour. To make the micropatterns fluorescent, 0.3 µg of CNA35 protein labelled with mCherry fluorescent protein was added to the final solution to nonspecifically bind to the fibronectin.

### Cell Culture

hTERT-immortalized Retinal Pigment Epithelial cells stably expressing LifeAct GFP (RPE1-hTERT LA GFP) were generated using lentriviral transduction. They were cultured in Dulbecco’s Modified Eagle Medium/F-12 with L-Glutamine (DMEM-F12, Capricorn scientific), supplemented with 10 % Fetal Bovine Serum (FBS, Sigma Aldrich), and maintained at 37 ^*°*^C and 5 % *CO*_2_. It was important not to let the cells reach 100 % confluency before splitting, as this prevented them from spreading completely on the patterns. At this step, the cells were used either for immunostaining or microrheology experiments.

#### Preparing cells for immunostaining

Cells were detached by incubating with Trypsin-EDTA (Sigma Aldrich, diluted from 0.25 to 0.05 %) at 37 ^*°*^C for less than 3 min. Next, they were resuspended in media, seeded on the micropatterns and kept at 37 ^*°*^C and 5 % *CO*_2_ for 1 hour.

They were then washed three times with Dulbecco’s PBS containing *MgCl*_2_ and *CaCl*_2_ (DPBS, Sigma Aldrich), by pipetting the buffer up and down very gently to prevent the cells from peeling off the surface. Most of the DPBS was gently removed from the coverslips, and the cells were fixed by 4 % paraformaldehyde (PFA). After 10 minutes, PFA was gently removed and the coverslips were washed 3 times with DPBS. During each wash step, PBS was added to the coverslips and incubated for 5 minutes before removal. The DPBS was then removed and the coverslips were incubated in blocking solution for 1 hour. Next, the samples were incubated with primary antibody for 2 hours and subsequently washed 3 times with PBS, following the same washing procedure described above. The cells were then incubated with secondary antibody for 45 min and washed a final three times with PBS. The blocking solution was prepared by supplementing PBS with 10 % Goat serum (Gibco) and 0.2 % Triton x 100 (Carl ROTH). The primary antibodies were diluted in the blocking solution as follows: Golgin-97 mouse IgG1 Mococlonal CDF4 (Invitrogen, 1:100 dilution) for the Golgi apparatus; SERCA2 ATPase Monoclonal IID8 (Invitrogen, 1:500 dilution) for the endoplasmic reticulum; and Alpha Tubulin Monoclonal (Invitrogen, 1:100 dilution) for microtubules. For the secondary antibody, Goat anti-mouse IgG1 Cross-Adsorbed Secondary Antibody Alexa Fluor 647 (Invitrogen) was diluted 1:500 in blocking solution. To stain the nuclei, 20 mM Hoechst 33342 (Thermo Scientific) was added to the secondary antibody solution at 1:1000 dilution.

#### Preparing cells for Microrheology

Cells were detached as described for immunostaining. After detachment, they were resuspended in media supplemented with a 1:1000 dilution of 1 µm blue fluorescent Carboxyl Polystyrene beads (Bangs Labs), 1 µM SiR-DNA (Spirochrome), and 10 µM Verapamil (Spirochrome). The cells were then seeded on the micropatterns and maintained at 37 ^*°*^C and 5 % *CO*_2_ for 45 min. Next, the micropatterns were washed by pipetting media 10 times to remove non-adhered cells and excess beads. A double-sided tape (DST1950, Thorlabs, United States) was cut into a rectangular frame (10 mm ×10 mm inner frame size) and stuck to a clean microscope slide. The inner space was filled with media supplemented with 10 mM HEPES, alongside SiR-DNA and Verapamil at the same concentration used previously. Finally, the coverslips were attached to the tape and the resulting chamber was sealed with silicon vacuum grease.

### Viscoelastic Hydrogel

The viscoelastic hydrogel is polyacrylamide-based with a dextran component to increase viscosity. The premix includes 147.6 µL of Acrylamide solution (Sigma-Aldrich, A4058-100ML, already 40 %w*/*v), 18.45 µL of N,N’-Methylethylenebisacrylamide solution (Sigma-Aldrich, M1533-25ML, already 2 %(w*/*v)) and 0.886 µL of Acrylic Acid (Sigma-Aldrich, 14230-100G). 2.14 g of Dextran solution (Sigma-Aldrich, 31389-25G, diluted 10 %w*/*v in water) was prepared. 1 µm diameter microspheres (Polysciences, Inc., Fluoresbrite Carboxy NYO 1.0 Micron Microsphere, 18449) were diluted 1:100 in water. As a catalyst N,N,N’,N’-Tetramethylethylenediamine (Sigma Life Science, T7024-25ML) was used. The reaction starter was Ammonium Persulfate (Sigma-Aldrich, A3678-25G) that was diluted 10 mg:100 µL in water just before the experiment. For an experiment 79.5 µL premix, 656 µL Dextran solution, 15 µL bead solution, 7.5 µL APS solution and 2.25 µL TEMED were mixed and quickly put into the sample (35 µL for microrheology, 603 µL) for macrorheology). The sample was measured after *t* ≥24 h to ensure complete polymerization which was checked using a conventional rheometer (Anton Paar Physica MCR 501) with an Anton Paar CP50-1 cone plate.

### Actin Reconstituted Network

Actin reconstituted networks were prepared in four steps

#### Bead passivation

Fluorescent carboxylated polystyrene beads (1 µm diameter, 1 %(w*/*v), FC04F, Bangs Laboratories, Fishers, IN, USA) were passivated with *α*-methoxy-*ω*-amino polyethylene glycol (CH_3_O −PEG −NH_2_, 122000-2, Rapp Polymere, Tübingen, Germany) to prevent non-specific protein adsorption to the beads, following established protocols [26, 42]. Briefly, the beads were washed twice in 50 mM 2-(N-morpholino)ethanesulfonic acid (MES; 41320078-1, BioWORLD, Dublin, OH, USA), pH 6. The carboxyl groups present on the bead surface were then activated through a 1-ethyl-3-(3-dimethylaminopropyl)carbodiimide (EDC; 22980, Thermo Fischer Scientific, Waltham, MA, USA) N-hydroxysuccinimide (NHS; 24500, Thermo Fischer Scientific, Waltham, MA, USA) (EDC-NHS) reaction performed at 37 ^*°*^C and 600 rpm on a shaker (ThermoMixer C, Eppendorf SE, Hamburg, Germany) for 15 min. Any residual products from the activation reaction were removed via a two washing cycles using 50 mM MES buffer. Then, the beads were resuspended in borate buffer (0.05 M, pH 8.5) (bioPlus, 40220048, BioWORLD, Dublin, OH, USA). Then, PEGylation was performed adding 0.297 M CH_3_O–PEG–NH_2_ to the beads and incubation for 150 min at 37 ^*°*^C and 600 rpm. Finally, the beads were extensively washed to remove residuals from the passivation and resuspended in 50 mM MES buffer to a final concentration of 10 mg mL^*−*1^ and stored at 4 ^*°*^C.

#### Actin purification

Actin was purified from rabbit skeletal muscle acetone powder, snap frozen and stored in G-buffer (5 mM Tris, pH 8.0; 0.2 mM Na-ATP; 0.2 mM CaCl_2_; 0.5 mM DTT) at −80 ^*°*^C as previously described [43, 44]. Before further use in the experiments, the concentration of unlabeled actin was measured using a UV-vis spectrophotometer (NP80, Implen GmbH, München, Germany) at 290 nm.

#### Preparation of actin network samples for microrheology

10 µL of passivated beads in MES buffer were centrifuged at 9000 ×*g* for 1× min and resuspended in 25 µL of 1× G-buffer and sonicated for 5 min. Unlabeled G-actin at a concentration of 95.2 µM was first incubated on ice for 2 h to reduce aggregation and ensure longer actin filaments in the networks. The actin polymerization mix contained 9.6 µM, 5.0 µM or 2.9 µM for a mesh size of 400 nm, 600 nm or 800 nm [45], respectively, as well as 20 mM Tris (pH 7.5), 4 mM MgCl_2_, 100 mM KCl, 3 mM ATP, 0.25 %(w*/*v) BSA, and 0.4 mg mL^*−*1^ passivated beads, and an oxygen scavenging system including 16 mM DTT, 2.5 mg mL^*−*1^ glucose, 83 µg mL^*−*1^ glucose oxidase, 33.3 µg mL^*−*1^ catalase. The polymerization mixture was incubated at room temperature for 5 min to promote actin polymerization. Subsequently, Alexa Fluor 647-conjugated phalloidin (Thermo Fisher Scientific, USA) was added at a molar ratio of 1:2 to label and stabilize the actin filaments.

#### Preparation of microrheology chambers

Microrheology chambers were prepared following previously published protocol [42]. Briefly, 1 mm-thick microscopy slides (24 mm ×60 mm; VWR International, Radnor, PA, USA) and no. 1.5 coverslips (24 mm × 24 mm; VWR International) were cleaned with 2-propanol, sonicated for 45 min and dried with nitrogen gas. Double-sided tape (tesafix 4959, Tesa, Norderstedt, Germany) with 100 to 120 µm thickness was cut into a rectangular frame (9 mm×8 mm inner frame size) and placed on the cleaned microscope slide. 8 µL of actin network solution were placed in the center of the frame. A hydrophobic rectangular border was drawn on the clean coverslip using a 2 mm PAP pen (Sigma-Aldrich, St. Louis, MO, USA). The coverslip was then used to close the chamber. Finally, the chamber was sealed using vacuum grease.

### Optical Setup

An optical tweezers was used to perform active and passive microrheology experiments. It was coupled to a Nikon Ti2 inverted microscope equipped with a Yokogawa CSU-W1 spinning-disk confocal unit. To trap the beads, a 5 W, 1064 nm laser (YLR-5-LP, IPG Photonics) was expanded to a beam waist of 5 mm and focused using 1.2 N.A. objective (CFI Plan Apo VC 60XC WI, Nikon Corporation), where its intensity was adjusted to provide 60 mW at sample position. The laser beam could be steered in both x and y directions with precision of 1.7 nm at the sample position using an xy acousto-optic deflector (DTSX-400-1064, AA Opto-Electronic) controlled by a variable-frequency driver (voltage controlled oscillator, DRFA10Y2X-D41k-34-50.110, AA Opto-Electronic). Most of the laser light was subsequently captured by a force sensor module (Lunam T-40i; Impetux Optics), which projected the back focal plane of the condenser to a Position-sensitive detector (PSD) to directly quantify the forces [46].

To measure the position of the bead with high precision, a 976 nm, 500 mW laser was utilized (BL976-PAG500, Thorlabs) and the intensity was adjusted to 1 mW to avoid forming an effective secondary trap. It was collected by the force unit, but the back-focal plane of the condenser was projected to another PSD. Before each round of microrheology, the voltage to position conversion factor was calibrated by moving the bead through the laser beam using a piezo stage (P-545.3R8H PInano XYZ, Physik Instrumente). The position detection was with nm precision.

### Microrheology

#### Active Microrheology

Mechanical properties of the system can be quantified using active microrheology (SI Fig. 3). In this type of measurement, the bead was trapped and oscillated by a specific frequency *f* in either x or y direction, where the oscillation amplitude of the laser was 200 nm. Both trap force *F* (*t*) and bead position *x*(*t*) were collected using the position sensitive detectors. These quantities are related to each other via a convolution relation

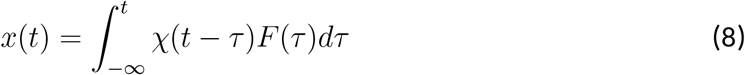

where *χ*(*t*) called response function, quantifies how the environment responds to the applied deformations. By using the convolution theorem, Eq. 8 simplifies to

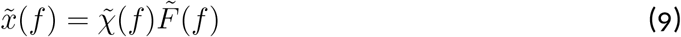

where 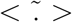 denotes Fourier transform. Therefore, by taking Fourier transform of measured *F* (*t*) and *x*(*t*) and using Eq. 9, the response function could be quantified. Then by using the generalized Stokes-Einstein equation [47, 48], the mechanical properties of the material could be calculated as follows:

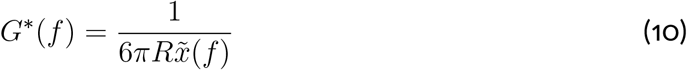

where *G*^*∗*^(*f*) is called complex shear modulus and *R* is the bead radius. The real part *G*^*′*^(*f*) and imaginary part *G*^*′′*^(*f*) of *G*^*∗*^(*f*) are called storage and loss modulus, respectively and quantify the elastic and viscous behavior of the material.

#### Passive Microrheology

The non-equilibrium activity of the system can be quantified using passive microrheology (SI Fig. 3). In this type of measurement, the spontaneous fluctuations of a non-trapped bead were captured using the position detection laser. To achieve this, position of the bead *x*(*t*) was recorded for 5 consecutive times for a duration of 5 s per tracking window. Subsequently, the average power spectral density (PSD) of the fluctuations 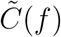, was calculated as follows

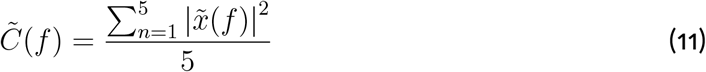

Assuming the system is in thermodynamic equilibrium, the PSD can be related to imaginary part of response function *χ*^*′′*^(*f*) via the Fluctuation-Dissipation Theorem (FDT) [49].

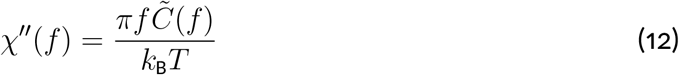

However, living cells are inherently non-equilibrium systems, and therefore Eq. 12 is violated. In such active systems, the magnitude of the fluctuations captured by passive microrheology is higher than what would be expected under thermal equilibrium. To account for this, an effective energy can be defined as *E*_eff_(*f*) = *E*_active_(*f*) + *k*_B_*T* to modify FDT as follows

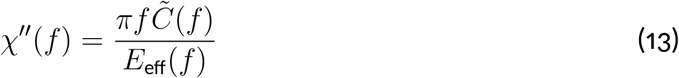

therefore, by independently measuring 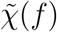 via active microrheology and 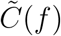 via passive microrheology, and substituting these values into Eq. 13, the active energy of the system can be estimated as

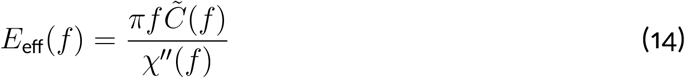

where any deviation from *k*_B_*T* is a hallmark of non-equilibrium activity.

### Microscopy

To reconstruct the average cell, different intracellular structures including the Golgi apparatus, endoplasmic reticulum, and microtubules, were immunostained in a series of independent experiments. Actin filaments, nuclei, and the underlying micropatterns were visualized across all datasets to serve as structural references. Z-stacks of single cells that were completely spread on the micropatterns were captured automatically using a custom, home-built LabVIEW program. To resolve the finer structural details, the raw fluorescence images were deconvolved using the Richardson-Lucy algorithm with Total Variation regularization [50] with *N* = 20 iterations and the regularization parameter *λ* = 0.001. The deconvolution was implemented on GPUs using cupy [51] for accelerated computation. The point spread function (PSF) used for deconvolution was measured by averaging over z-stacks of 302 green (excited by 488 nm laser) and 84 and red (excited by 560 nm laser) 100 nm fluorescent microspheres. Due to spectral proximity, the green PSF was used to deconvolved the blue (excited by 405 nm laser), while the red PSF was used for the far-red channel (excited by 640 nm laser). From the deconvolved stacks, maximum intensity projections were calculated to compress the three-dimensional structural data into a single two-dimensional plane. To remove rotational and translational degrees of freedom across the dataset, the fluorescent micropattern in each image was used as a fixed reference frame to globally register the images. To prevent background fluorescence from introducing structural artifacts, the images were segmented using a fine-tuned version of *µ*SAM, which is built upon the Segment Anything Model [52, 53]. All pixel intensities outside the segmented cell boundary were set to zero. Following segmentation, the remaining image intensities were normalized to integrate to unity. This mathematical normalization allowed each individual image of a specific structure, denoted as *Im*_*i*_, to be treated formally as a two-dimensional probability density function *µ*_*i*_ ∈ *P* (ℝ^2^). Converting the images into normalized probability distributions was a requirement for computing the average cell using optimal transport.

### Optimal Transport and Wasserstein Barycenter

To construct a geometrically faithful average cell from a population of *N* cell images {Im_1_, Im_2_, … , Im_*N*_ *}*, standard pixel-wise arithmetic averaging is insufficient as it results in unphysical blurring. Instead we use the Wasserstein geometry induced by optimal transport to compute a more meaningful notion of average. For this we normalize the total intensities in each image to one and interpret images as continuous probability distributions *µ*_*i*_ ∈ *P* (ℝ^2^), where *x* = (*x*_1_, *x*_2_) is the 2D pixel position. There are several mathematically equivalent ways to formulate optimal transport problems and we refer to [29] for an approachable introduction. Numerically we use the Python package `MultiScaleOT`which uses the Sinkhorn algorithm to solve an entropy regularized variant of the Kantorovich formulation.

Conceptually, our methodology is best explained by using a fluid dynamics formulation [54]. To compute *W*_2_(*µ*_*i*_, *µ*_*j*_) we warp the source image continuously into the target image like a moving fluid over a time interval *t* ∈ [0, 1]. Let *µ*(*t, x*) describe the evolving, time-dependent image intensity density at time *t* and coordinate *x* ∈ ℝ^2^. The movement of infinitesimal mass particles is determined by a spatio-temporal vector field *v*(*t, x*) ∈ℝ^2^ that represents the velocity of mass particles at that specific time and position. The effect of the velocity field on the image intensity is described by the continuity equation, *∂*_*t*_*µ* + ∇ · (*µv*) = 0, which encodes conservation of mass. The squared 2-Wasserstein distance is then defined as the minimum action required to implement the image transformation

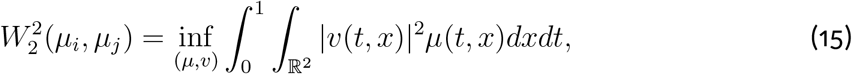

subject to the mass conserving continuity equation and the temporal boundary conditions *µ*(0, *x*) = *µ*_*i*_(*x*) and *µ*(1, *x*) = *µ*_*j*_(*x*). The inner integral represents the physical kinetic energy of the mass particles at a given instant. Therefore, the 2-Wasserstein distance is given by the minimum average kinetic energy required to continuously morph one image configuration into another. It can be shown that for minimizers of (15) the particles move with constant speed along straight lines from their initial to their final positions (Fig. 2). Consequently, 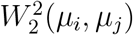 also equals the minimum average squared distance that each mass particle must be moved to rearrange *µ*_*i*_ into *µ*_*j*_.

*W*_2_ describes a mathematical distance on the set of probability densities. In particular, it satisfies the triangle inequality and the minimizing interpolation *t* → *µ*(*t*, ·) describes a shortest path with respect to this distance, connecting the initial and final distributions. This provides a natural way to interpolate smoothly between two densities.

More generally, one can also define a metric average of more than two images with respect to the distance *W*_2_, the so-called *Wasserstein barycenter µ*_*b*_ [55]. It is given as the unique minimizer of the average squared distances to all image intensities in the dataset:

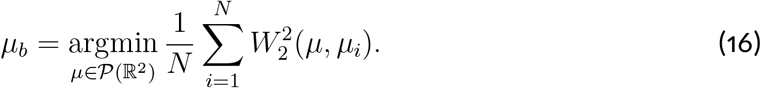

Mathematically, the fluid mechanics formulation (15) induces a formal abstract Riemannian manifold, where each individual probability distribution *µ*_*i*_ corresponds to a single distinct point on this geometric surface [56] (SI Fig. 8). At the location of the barycenter *µ*_*b*_ on this curved manifold, we can attach a flat tangent space. Every sample image *µ*_*i*_ can be projected from the curved manifold onto this flat tangent space where it is represented by a static velocity vector field *v*_*i*_(*x*) which is given by the optimal initial (i.e. at *t* = 0) velocity field in the optimal interpolation from *µ*_*b*_ to *µ*_*i*_ according to (15). One can then approximate *W*_2_(*µ*_*i*_, *µ*_*j*_) from the discrepancy between the velocity fields *v*_*i*_ and *v*_*j*_, weighted by *µ*_*b*_, as follows:

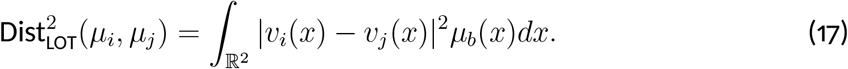

This is called the linearized optimal transport (LOT) approximation [56, 57]. One advantage is that instead of having to solve the optimization problem for *W* (*µ*_*i*_, *µ*_*j*_) for all *O*(*N* ^2^) pairs of samples, instead one must only solve *N* problems to compute the initial velocities *v*_*i*_ and then obtain all pairwise approximate distances much faster. One can also use this to compute linear averages of velocity fields or even apply PCA to a dataset of velocity fields to extract orthogonal dominant modes of deformation. In particular, this approximation can also be applied directly to (16) to obtain an approximation of *µ*_*b*_, which is challenging for large *N* . The barycenter itself is mapped to the origin of this tangent space (*v*_*b*_ = 0) and indeed the distance *W*_2_(*µ*_*i*_, *µ*_*b*_) of any sample *µ*_*i*_ to the barycenter *µ*_*b*_ is exactly given by

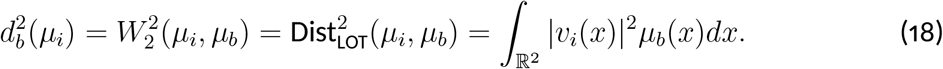

We handled numerical computations involving linearized optimal transport with the Python package `LinOT`. As demonstrated by Fig. 2, on our dataset the linear structure on velocity fields, induced by optimal transport, is much more appropriate than the original linear structure on the image intensities 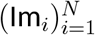.

### Gaussian Process Transformation

A direct aggregation of data points does not result in a template mappable to the *W*-cell template. In fact, although confining the cells onto micropatterns standardizes the global morphology of them, the internal organization specially the nucleus position remains variable. This variability introduces a significant artifact if raw coordinate data is used: a bead physically close to the real nucleus in an individual cell might appear far from the nucleus when overlaid onto an W-cell template, and vice versa.

To eliminate this spatial bias and ensure that relative physical distances are preserved, the coordinates of each individual cell space were transformed into a standardized *W*-cell space using Gaussian Process interpolation [58, 59]. To achieve this, the joint boundary coordinates of both the actin cortex and the nucleus were extracted for each individual cell. This generated a combined source boundary set, 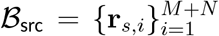, where **r**_*i*_ = [*x*_*s,i*,_*y*_*s,i*_] represents the *i*^th^ sampled point on either the actin or nuclear boundary, and *M* and *N* denote the total number of sampled points along the actin and nuclear boundaries, respectively. This source set was paired with a corresponding target boundary set,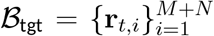 , where **r**_*t,i*_ = [*x*_*t,i*_, *y*_*t,i*_] is the coordinate on the W-cell template corresponding to the *i*^th^ source point (SI Fig. 9).

For the actin boundary, **r**_*t,i*_ was defined as the nearest point on the *W*-actin template relative to **r**_*i*_, since the cell periphery exhibited only minor deviations from the micropattern shape. However, this proximity-based logic was insufficient for the highly variable nuclear positions. To overcome this limitation, optimal transport was utilized to calculate displacement vectors, which were then used to systematically assign the corresponding target points on the template.

Using these boundary mappings, any individual bead position **r**_ind_ within the cytoplasmic region was smoothly transformed into a corresponding position **r**_avg_ in the W-cell frame via:

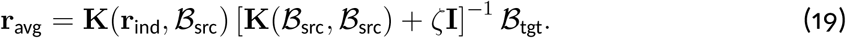

Here, **K**(ℬ_src_, *ℬ*_src_) is an (*M* + *N*) × (*M* + *N*) covariance matrix evaluating the kernel function *C*(**r**_*a*_, **r**_*b*_), between all pairs of source boundary points, while **K**(**r**_ind_, *ℬ*_src_) is a 1× (*M* + *N*) row vector evaluating the kernel between the specific bead position and all source boundary points. The kernel was defined as

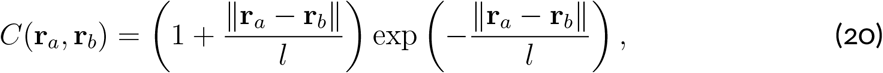

where the characteristic length scale was set to a large value of *l* = 65 *µ*m to prevent local grid compression or unphysical warping of the space. The term *ζ***I** in Eq. 19 is a regularization matrix used to ensure numerical stability during inversion, with *ζ* = 0.01. Transforming coordinates through this framework guarantees that beads originally adjacent to the nucleus remain adjacent to the W-nucleus, while distant beads remain distant.

### Nested Cross-validation for Spline Model Evaluation

We modeled spatial dependence of the rheological parameters using a two-dimensional generalized additive model (GAM) [60] with a tensor-product spline basis in the spatial coordinates *x* and *y*. Each marginal coordinate was represented by a cubic B-spline basis with 15 basis functions, which were combined into a two-dimensional tensor-product basis. The spatial domain of the basis was fixed from the full coordinate range, including the cell boundary, with an additional 2% padding at each edge. Moreover, since the micropattern geometry is symmetric along the vertical axis, which is also reflected in the average localization of cytoplasmic structures, we assumed that the mechanical properties of the cytoplasm are likewise symmetric along the same axis. Therefore, to enforce this symmetry and maximize the statistics, reflection symmetry about *x* = 0 was imposed on the data level during fitting by augmenting each training set according to

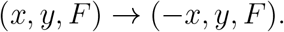

Here, *F* is any one of the rheological parameters considered. We applied this augmentation only after defining the respective training subset, so we never introduced validation and test observations into the training data through the symmetry operation.

The flexibility of the spline is controlled by the GAM regularization parameter *λ*. We applied a common smoothing parameter along both spatial dimensions. Increasing *λ* increases the penalty on spatial curvature and therefore favors smoother surfaces, whereas decreasing *λ* allows progressively more spatial variation. To tune the degree of smoothing, *λ* was selected from 25 logarithmically spaced values between 10^*−*5^ and 10^3^ by cross-validation.

To separate model tuning from the final evaluation of predictive performance, we used nested cross-validation [61, 62]. We first randomly divided the complete dataset into 15 outer folds. In each outer iteration, one fold was held out completely as the test set, while the remaining observations formed the outer training set. The held-out test data were not used during either spline fitting or selection of *λ*.

Hyperparameter selection of *λ* was performed exclusively within the outer training set using an additional 30-fold inner cross-validation. For each candidate value of *λ*, the GAM was fitted to the corresponding inner training folds and evaluated on the inner validation fold. Predictive performance was quantified by the mean squared error,

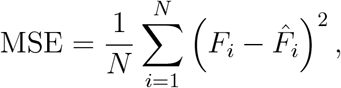

where *F*_*i*_ is the *i*th data point’s value of the mechanical parameter considered, and 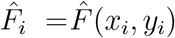 is the corresponding model prediction. The value of *λ* minimizing the mean inner-validation MSE was selected independently for each outer fold. A new GAM was then fitted to the complete outer training set using this selected value of *λ* and evaluated on the previously untouched outer test set. Thus, every observation was used once as out-of-sample test data and was not involved in selecting the smoothing parameter used to predict it.

We compared the spatial GAM with a spatially invariant constant model. For each outer fold, the constant model was fitted using only the outer training set, with its predicted value defined as the arithmetic mean

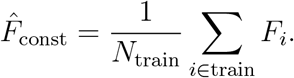

We then used this single value to predict every observation in the corresponding outer test set. The GAM and constant model were therefore evaluated on the same unseen observations, so their difference in predictive performance directly measures whether spatial position contributes predictive information beyond the global mean.

For each outer fold, we defined the improvement of the spatial model as

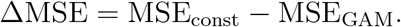

Positive values of ΔMSE indicate improved out-of-sample prediction by the spatial GAM, whereas values around zero indicate no predictive advantage over the spatially invariant baseline. For each channel, we perform a one-sided paired *t*-test across the outer folds to test for ΔMSE *>* 0, and correct the resulting *p*-values for multiple comparisons (Bonferroni) across all twelve tests, i.e. six channels evaluated with and without cell-specific biases (see below). The results are shown in SI Fig. 10. None of the channels shows a significant improvement of the GAM over the respective baseline, and we therefore find no evidence for discernible spatial structure in the present data. Although any spline we fitted is spatially modulated, the patterns can always be attributed to random noise. In SI Fig. 11, we show that spline patterns are not consistent across disjoint subsets of the data, matching our interpretation of the cross-validation analysis presented here.

As an additional control for the dataset’s hierarchical structure, we also considered spline models with cell-specific biases. Here, measurements from the same cell were treated as belonging to a common experimental group whose response could shift relative to the global population mean through a cell-specific offset parameter. This accounts for the possibility that individual cells exhibit systematic biases that are not captured by the spatial spline itself. Such cell-to-cell variation contributes to the dataset’s overall variance and may obscure spatial dependencies within individual cells. Accounting for these effects could improve within-cell prediction and increase the spline model’s sensitivity to spatial structure shared across cells. Fitting these models only collects gradient information from measurements within the same cell, and therefore only cells with at least 2 measurements were considered. We evaluated the models with cell-specific biases using the same cross-validation scheme as above. However, data points were only left out if the remaining cell groups still contained one or more measurements. This is required, as the prediction requires estimation of the cell bias. The prediction scoring was performed against the estimated cell average as baseline. The analysis of the models with cell-specific effects also did not reveal any evidence for detectable spatial structure in any of the channels (SI Fig. 10).

### Spatial Autocorrelation Functions

The autocorrelation function was computed from a stack of *N* images, {Im_1_, Im_2_, … , Im_*N*_ *}*, using the Wiener–Khinchin theorem, independently for each structure. Prior to analysis, each image was normalized to have a zero mean and unit variance (Im_*i*_ ← (Im_*i*_ −*µ*_*i*_)*/σ*_*i*_) to ensure that brightness or contrast differences between images do not bias the correlation results. According to this theorem, the 2D spatial autocorrelation function for an individual image Im_*i*_(*x, y*) is defined as

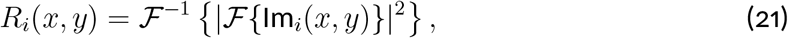

where ℱ denotes the 2D Fourier transform and (*x, y*) represent spatial coordinates. To obtain a 1D radial profile, each 2D correlation function *R*_*i*_(*x, y*) was radially binned at a 1-pixel resolution. The average value for the *n*-th radial bin is calculated as

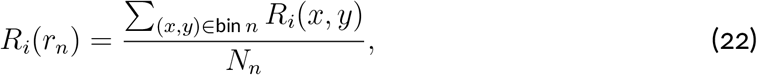

where *r*_*n*_ denotes the radial distance corresponding to the *n*-th bin and *N*_*n*_ is the number of pixels within that bin. Finally, the ensemble-averaged spatial correlation function 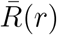 was obtained by averaging the profiles across all *N* images in the stack

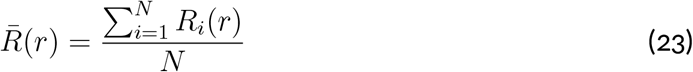

### Principal Component Analysis

To analyze mechanical fingerprint data (*p* ∈ {*A, B, α, β, E, γ* }), Principal Component Analysis (PCA) was performed. Prior to PCA, parameters spanning multiple orders of magnitude (*A, B*, and *E*) were log-transformed. They were then standardized using z-score normalization to ensure equal weighting across parameters with different physical units and scales:

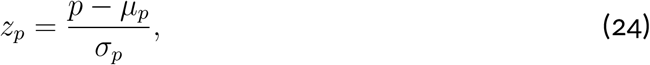

where *µ*_*p*_ and *σ*_*p*_ denote the mean and standard deviation of parameter *p*, respectively. The proportion of total dataset variance captured by each principal component *j* (explained variance) was calculated from the eigenvalues *λ*_*j*_ of the standardized covariance matrix:

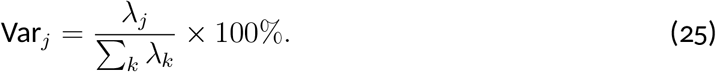

To determine how strongly individual parameters drive each principal component, the fractional parameter contributions were quantified using the squared loadings (*W* ^2^) across the first three principal components (*C*_1_, *C*_2_, *C*_3_):

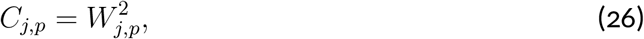

where *W*_*j,p*_ represents the eigenvector weight of the parameter *p* for the *j*-th principal component.

