## Supplemental Figures for "Structural order emerges without mechanical order inside living cells"

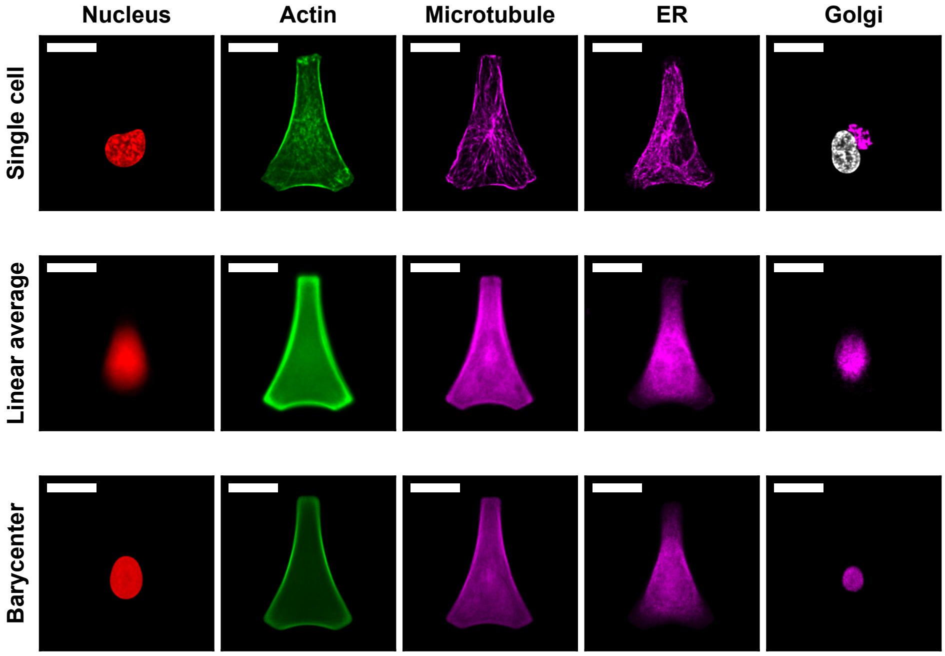


SI Figure 1: *Comparison between single-cell instances (first row), conventional linear averaging (second row), and Wasserstein barycenters (third row). Note that the representative single-cell panels display distinct cells, so the individual nuclear geometries vary across these examples. In the single-cell Golgi panel, the nucleus is overlaid in white to highlight the biological reality that the Golgi apparatus consistently localizes adjacent to the nuclear envelope. In contrast, both the linear averaging and Wasserstein barycenter methods fail to preserve this feature, resulting in an artifactual overlap where the averaged Golgi signal bleeds directly into the averaged nuclear domain.*

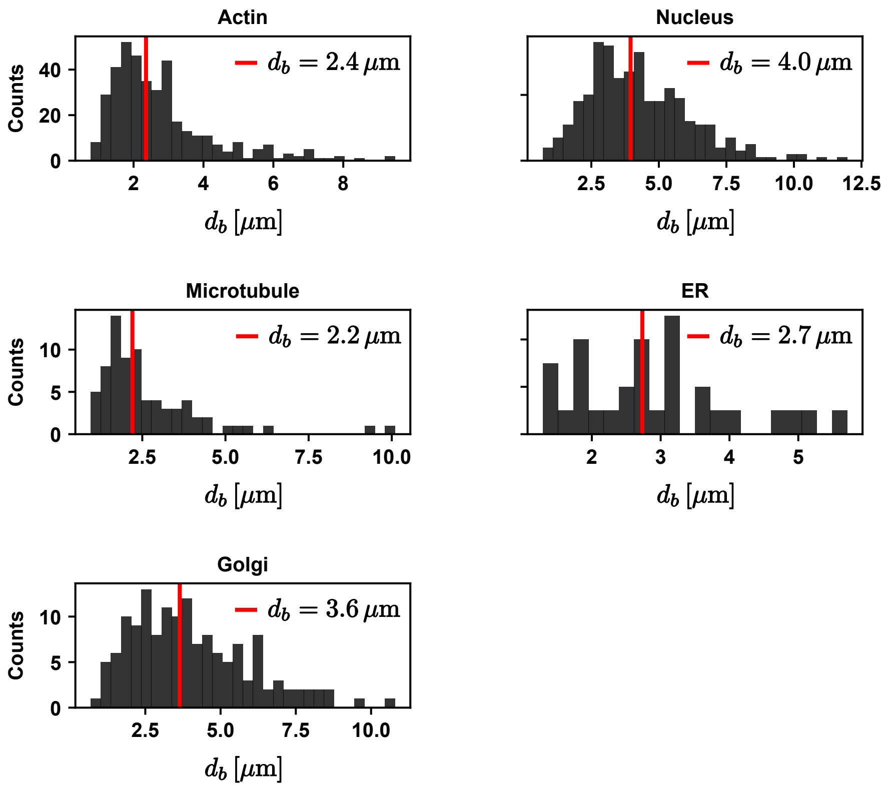


SI Figure 2: *Histogram of the linearized Wasserstein distance ($d_b$) for individual actin, nucleus, microtubule, endoplasmic reticulum, and Golgi apparatus images relative to their corresponding barycenter. Red lines show median values.*

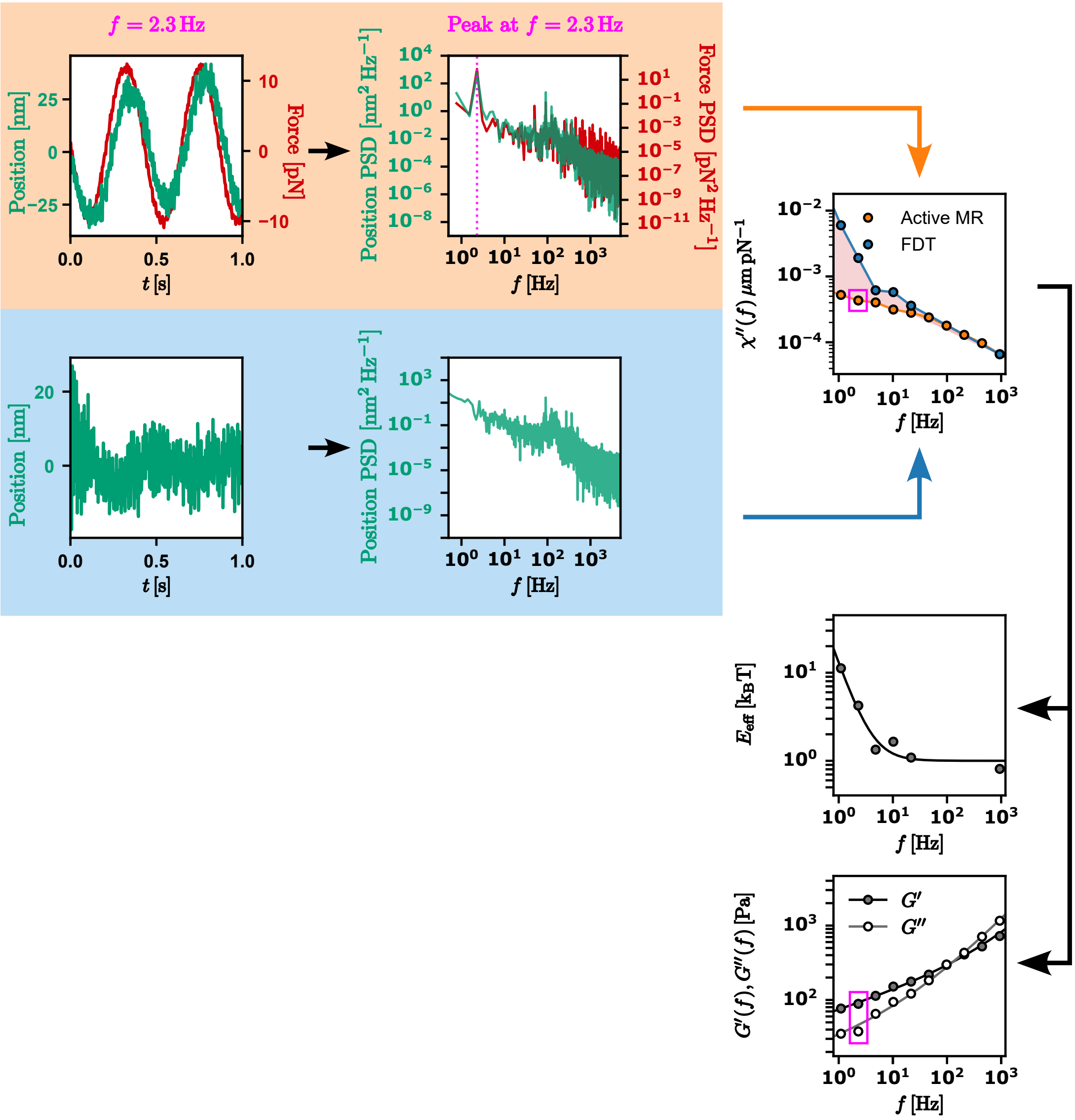


SI Figure 3: *Active (orange box) and passive (blue box) microrheology workflow for quantifying the response function, $\chi(f)$. The active microrheology pathway directly measures the complex response function $\chi(f)$, from which the complex shear modulus $G^*(f)$ is derived. Concurrently, the passive microrheology pathway utilizes the fluctuation-dissipation theorem (FDT) to estimate the dissipative imaginary component, $\chi''(f)$, from spontaneous thermal and active fluctuations. By comparing the active with passive dissipative $\chi''(f)$, the effective energy spectrum, $E_{\text{eff}}(f)$, can be quantified.*

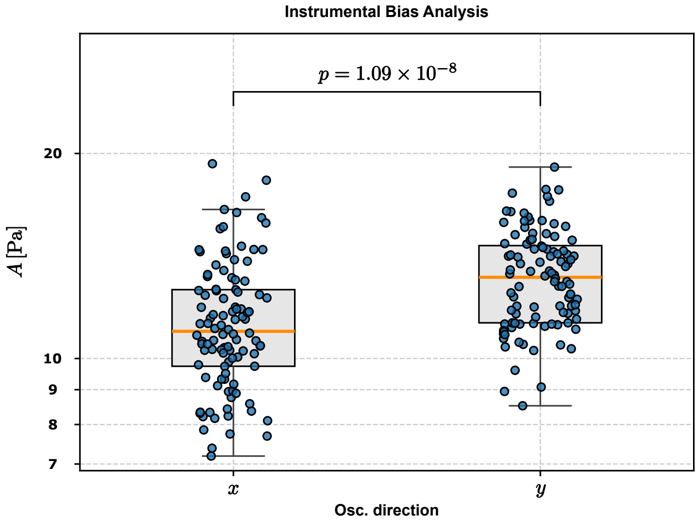


SI Figure 4: *Quantification of instrumental bias. Microrheology control measurements in an isotropic viscoelastic hydrogel reveal a $20\%$ systematic bias toward higher stiffness values ($A$) along the $y$-axis relative to the $x$-axis.*

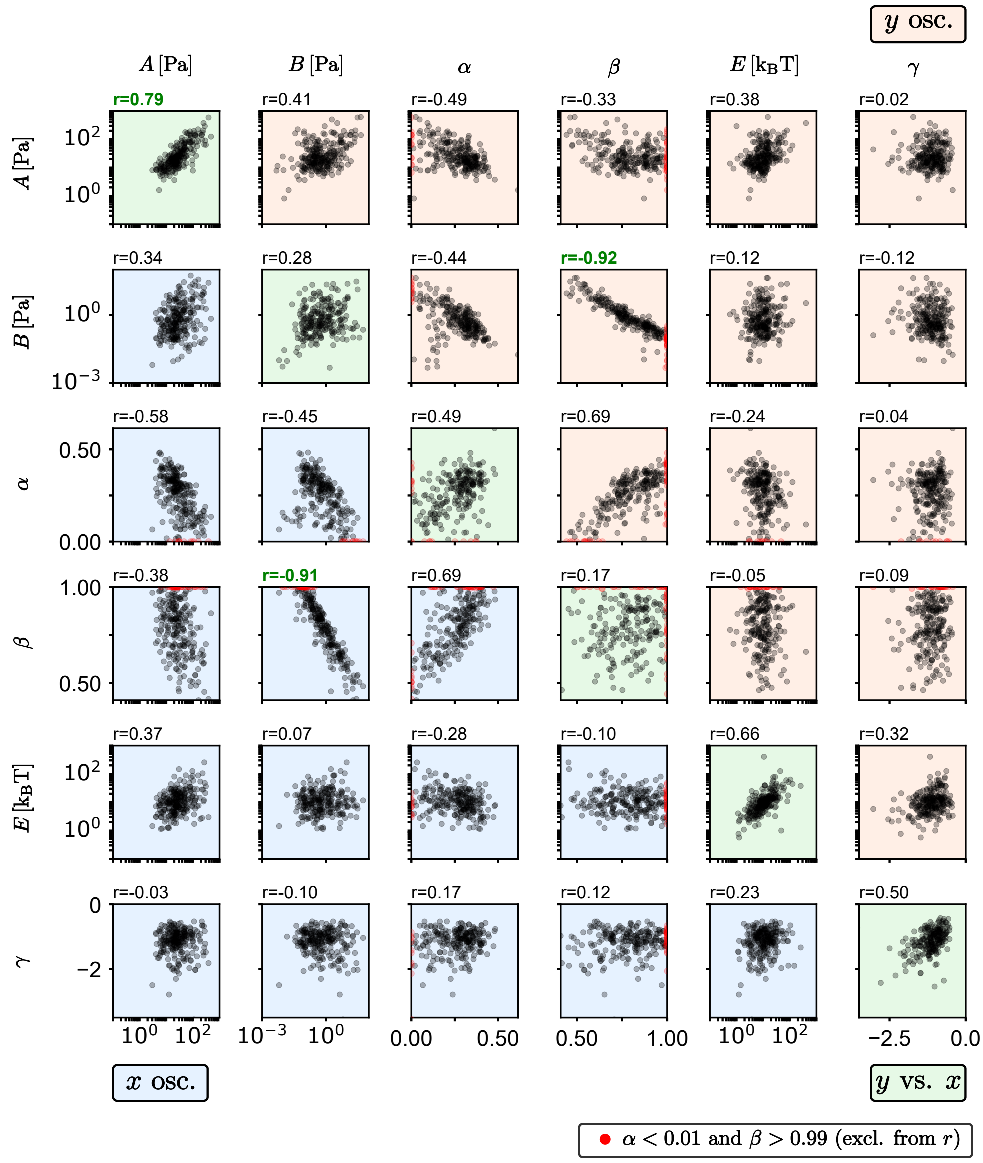


SI Figure 5: ***a**: Pairwise cross-correlation matrix of mechanical fingerprint parameters ($A$, $B$, $\alpha$, $\beta$, $E$, and $\gamma$). The lower triangle (blue background) displays parameter pairings measured along the $x$-axis, while the upper triangle (orange background) displays pairings along the $y$-axis. The Pearson correlation coefficient ($r$) is explicitly annotated for each pair of parameters. The diagonal shows parameter cross-correlation between $x$ and $y$ direction.*

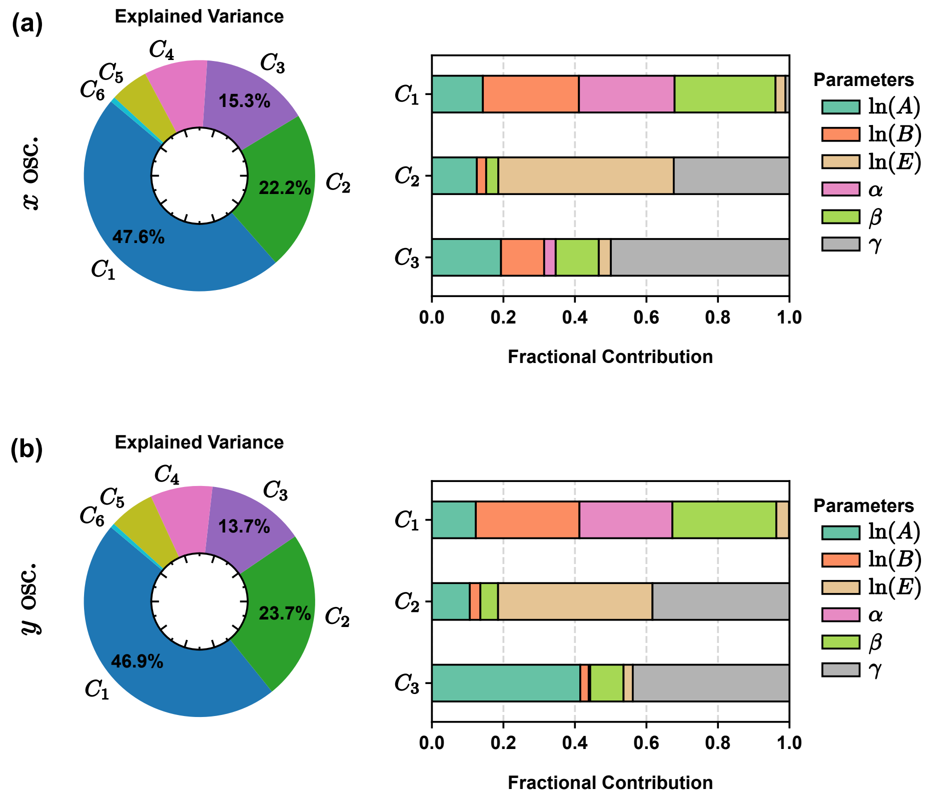


SI Figure 6: *Principal Component Analysis of the Intracellular Mechanical fingerprint.* ***a****: Results for the* $x$*-direction. The left panel shows the variance explained by all six principal components (*$C_1$*–*$C_6$*), with the first three components (*$C_1$*–*$C_3$*) accounting for nearly* $85\%$ *of the overall data variability. The right panel displays the contribution of each individual mechanical parameter to* $C_1$*,* $C_2$*, and* $C_3$*.* ***b****: similar to (a) but for* $y$ *direction.*

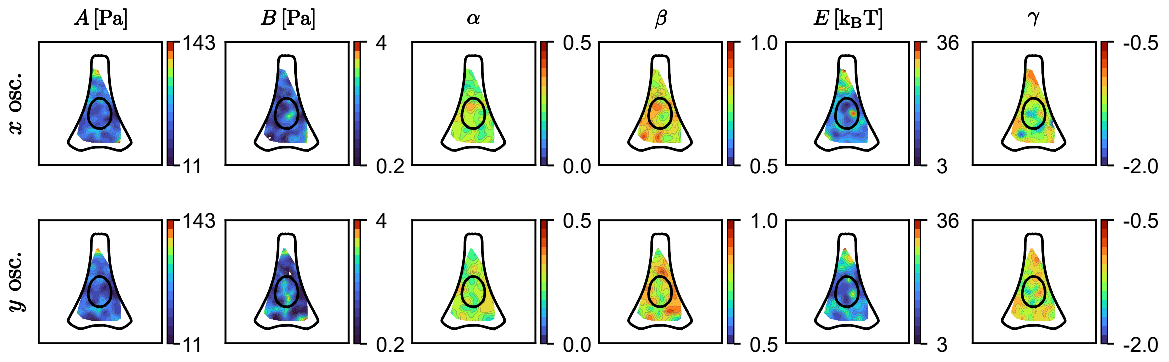


SI Figure 7: *Spatial mapping of mechanical fingerprint via Gaussian kernel regression. The spatial bandwidth is* $\sigma = 8\,\mu\text{m}$*, and* $k = 15$ *nearest neighbors per grid point are included. The log-normally distributed parameters* $A$*,* $B$*, and* $E$ *were* $\log$*-transformed prior to spatial interpolation. Contours are masked outside the domain defined by the intersection of the data convex hull and the outer actin boundary.*

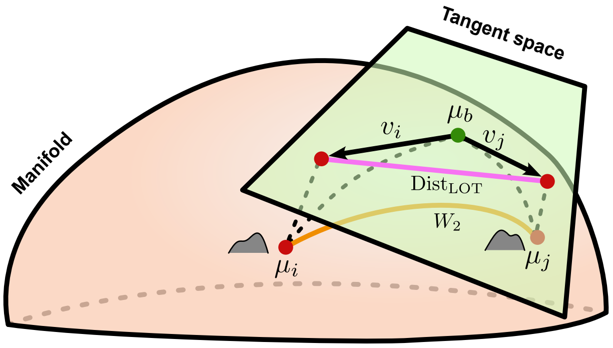


SI Figure 8: *Tangent space projection in Linearized Optimal Transport.Each probability distribution ($\mu_i$) corresponds to a point on a curved Riemannian manifold. By constructing a flat tangent space at the Wasserstein barycenter ($\mu_b$), each distribution $\mu_i$ can be projected onto this space and represented by its optimal initial velocity vector field $v_i(x)$. This linearization maps non-linear optimal transport distances to weighted Euclidean discrepancies between velocity fields, significantly simplifying pairwise distance computations.*

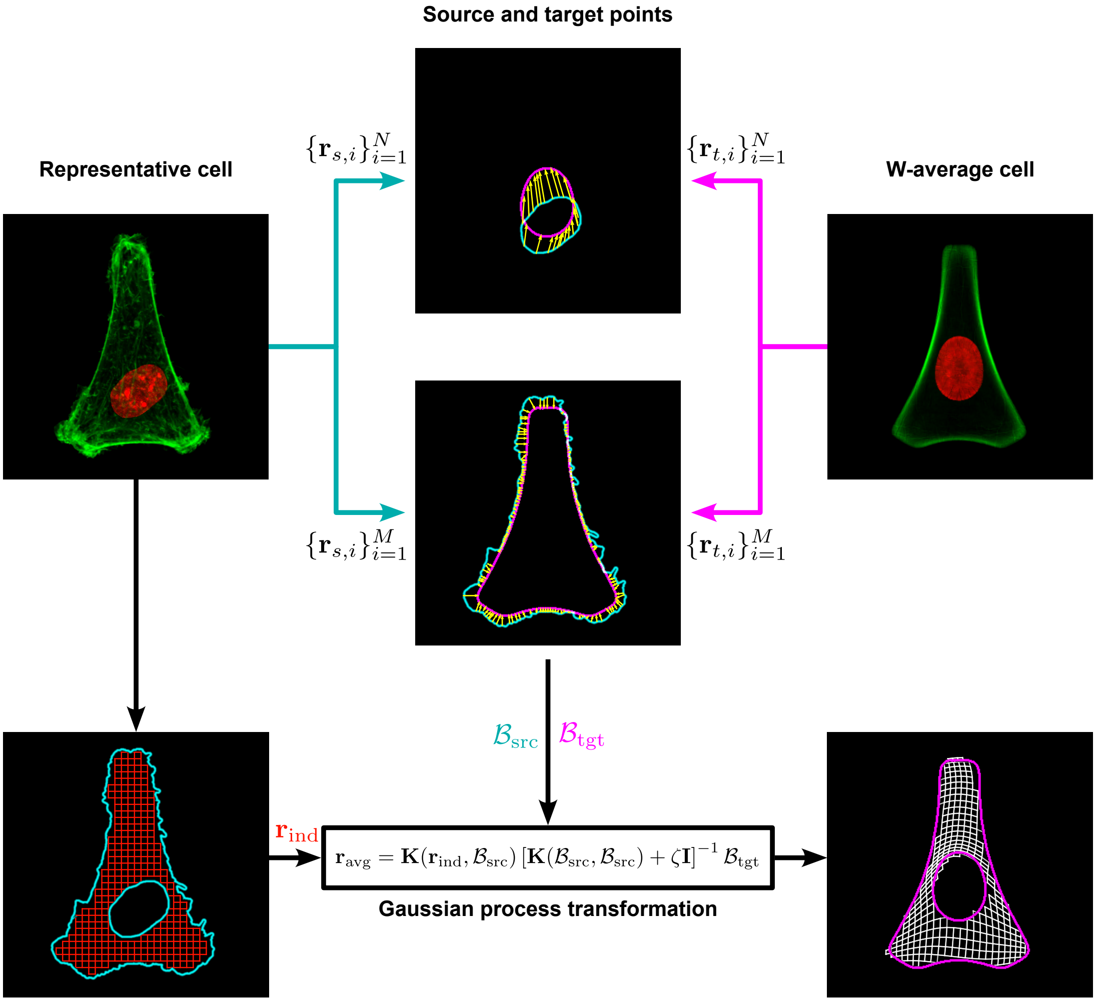


SI Figure 9: *Gaussian Process spatial alignment pipeline.Mapping of individual cytoplasmic microprobe coordinates ($\mathbf{r}_{ind}$) to the standardized $\mathcal{W}$-cell template ($\mathbf{r}_{avg}$). Boundary points from the cell periphery and nucleus ($\mathcal{B}_{src}$) are registered to target template boundaries ($\mathcal{B}_{tgt}$), driving a smooth Matérn-kernel Gaussian Process transformation that eliminates cell-to-cell spatial bias.*

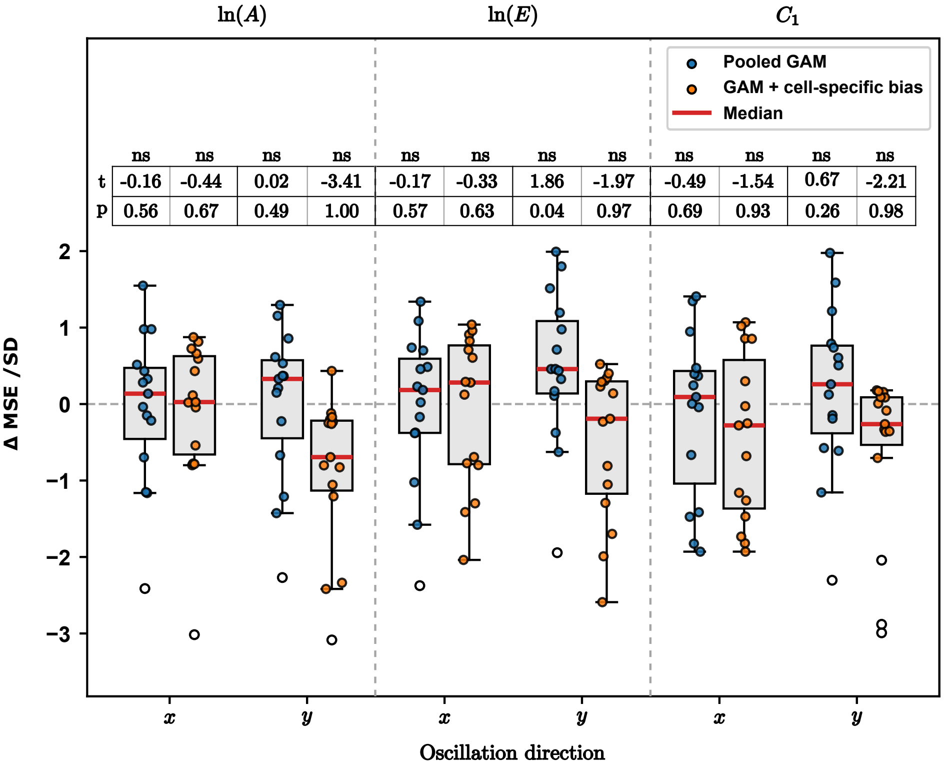


SI Figure 10: *Out-of-sample predictive performance of the spatial GAMs relative to their spatially invariant baselines, evaluated by nested cross-validation for* $\ln(A)$, $\ln(E)$ *and* $C_1$ *along the* $x$*- and* $y$*-directions. Blue: pooled GAM compared with the constant model,* $\Delta\text{MSE} = \text{MSE}_{\text{const}} - \text{MSE}_{\text{GAM}}$*. Orange: GAM with cell-specific biases compared with the cell-mean model,* $\Delta\text{MSE} = \text{MSE}_{\text{cell}} - \text{MSE}_{\text{GAM}}$*. Each point corresponds to one of the 15 outer folds, normalized by the standard deviation of the respective* $\Delta\text{MSE}$ *values; boxes indicate the interquartile range and red lines the median. Positive values reflect improved out-of-sample prediction by the spatial model, whereas values near zero indicate no predictive advantage over the respective baseline. For each box,* $\Delta\text{MSE} > 0$ *is tested using a one-sided* $t$*-test across the outer folds. The table above the boxes lists the* $t$ *statistic and the uncorrected* $p$*-value. To account for multiple comparisons, significance thresholds were Bonferroni-adjusted for all 12 tests, i.e.* $p < 0.05/12$ *(*),* $p < 0.01/12$ *(**) and* $p < 0.001/12$ *(***); ns: not significant. None of the tests reveals a significant improvement of the spatial model over its baseline.*

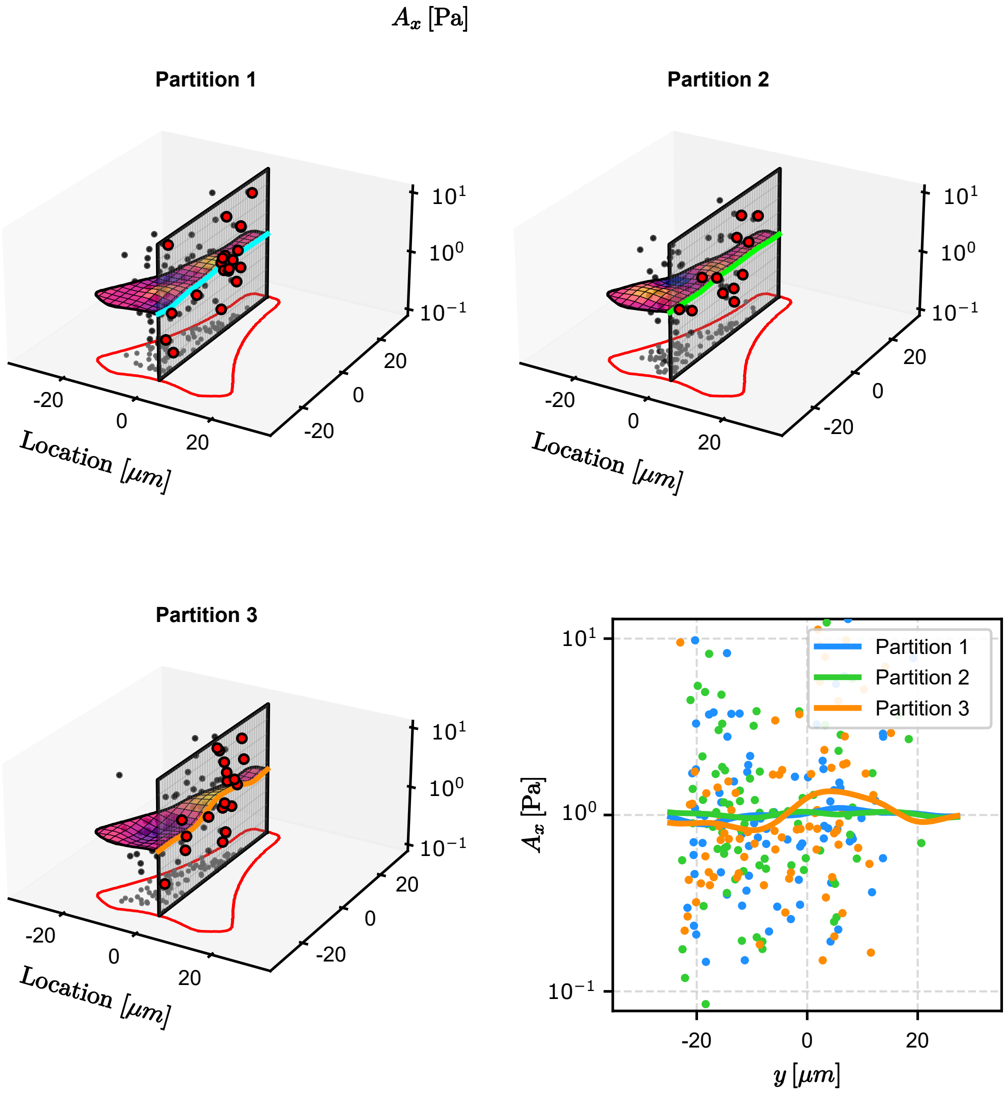


SI Figure 11: *Three separately fitted Generalized Additive Model (GAM) surfaces and underlying scatter data for parameter $A_x$, split across three disjoint data partitions. Cross-sectional profiles along the central symmetry plane ($x = 0\,\mu\mathrm{m}$) are highlighted in blue, green, and orange, with adjacent experimental data points (within a $\pm 2\,\mu\mathrm{m}$ tolerance) projected in red onto the slice plane. The lower-right panel displays the three cross-sectional profiles alongside all data points projected along the $x$-axis.The spatial variations of the splines show no consistent pattern across partitions. This illustrates that the apparent spatial features stem from the stochastic nature of the underlying data, rather than physically meaningful variation. The full joint dataset also shows no significant structure, as the nested cross-validation procedure described in the main text indicates.*
